# Label-free microscopy enables high-throughput identification of genes controlling biofilm development

**DOI:** 10.1101/2025.09.02.673883

**Authors:** M. R. Pratyush, Jojo A. Prentice, Rory A. Eutsey, Irina Mikheyeva, N. Luisa Hiller, Andrew A. Bridges

## Abstract

The biofilm mode of growth plays a critical role in microbial ecology and in the persistence of human pathogens. Yet, much remains unknown regarding the molecular determinants of biofilms in human pathogens. In this study, we present label-free analysis of biofilms (LFAB), an imaging approach that combines time-lapse, low-magnification brightfield microscopy with regional optical density measurements to quantify biofilm biomass. Unlike other approaches to biofilm biomass quantification, LFAB enables real-time, non-perturbative, and high-throughput monitoring of biofilms. We validated LFAB in diverse microbes and found that our measurements strongly correlate with traditional biofilm assays. We then used LFAB to identify and characterize critical factors mediating biofilm formation in *Streptococcus pneumoniae*, a major human pathogen whose biofilm lifecycle is known to be intimately related to colonization and infection. Initial characterization revealed that *S. pneumoniae* microcolonies form by radial expansion of attached cells, displaying reproducible morphology and growth dynamics. Screening of a transposon mutant library revealed that genes spanning carbohydrate metabolism, signaling, surface binding, cell wall synthesis, and adhesion impinge on the biofilm lifecycle of *S. pneumoniae*. We performed follow-up investigations of choline binding protein A (CbpA) and its adjacently encoded two-component system regulator, which we find are critical for the dynamics of microcolony biofilms in *S. pneumoniae*. Overall, this work establishes LFAB as a powerful approach for identifying and characterizing biofilm determinants across bacteria and uncovers key regulators of the biofilm lifecycle in a major human pathogen.

## Introduction

To colonize environmental and host niches, microbes often form surface-attached multicellular communities called biofilms.^1^ Biofilms are formed via the secretion of extracellular components that facilitate cell-to-cell and cell-to-surface adhesion.^2^ The biofilm matrix sterically restricts interactions between the cells and threatening agents such as bacteriophages and antibiotics.^3^ In addition, resident cells are protected from dislocation by fluid flow, and the high cell density that emerges from cell-to-cell attachments facilitates intercellular interactions such as the exchange of metabolites, cell-cell communication molecules, and genetic material.^1^ Biofilm forming bacteria are notorious for causing difficult-to-treat infections, as the biofilm lifestyle allows bacteria to evade the immune system and overcome clinical interventions.^4^ Thus, over the past several decades, the identification and characterization of molecular mechanisms and ecological principles underlying biofilm formation have become major areas of focus within microbiology. Ongoing efforts aim to manipulate the biofilm lifecycles of notorious pathogens as a novel strategy for treating recalcitrant infections.^5,6^

Given the ubiquity and importance of biofilms in medical and industrial settings, there is a broad interest in assessing the characteristics of biofilms formed by diverse microbial communities, species, and strains in a non-perturbative fashion. Furthermore, in genetically tractable organisms, there is an interest to identify the genes controlling biofilm development, which often requires high-throughput genetic screens. Existing methods of measuring biofilm characteristics are generally invasive, low throughput, and/or are unable to capture temporal dynamics. Perhaps the most widely used method to assess the propensity of a bacterial culture to form biofilms is the crystal violet (CV) assay.^7,8^ In this approach, bacteria are cultured, non-adherent cells are washed away, and remaining biofilm cells are stained using the CV dye. “Biofilm biomass” – a proxy for the number of biofilm cells in the community – can then be inferred by measuring the absorbance of solubilized CV using a spectrophotometer. While this approach has enabled countless discoveries and has been instrumental in the biofilm field, it imposes significant limitations: (1) the staining process kills cells, making real-time measurements impossible, (2) the numerous washing steps can dislodge biofilm structures, and (3) CV staining does not provide any information about biofilm morphology. Another common approach for measuring biofilm properties is confocal, light-sheet or super-resolution fluorescence microscopy, which, combined with advanced image analysis techniques, can yield reliable quantitative information on both biofilm dynamics and architecture.^9–14^ However, fluorescence-based imaging generally requires genetic manipulation of organisms of interest to introduce fluorescent probes, and samples are often subject to photodamage and photobleaching during image acquisition. The drawbacks of these and other assays for biofilm development have contributed to a diversity of approaches that are employed in biofilm literature, which in turn has led to inconsistent empirical standards for the objects that are qualified as biofilms.

Previous studies by members of our group began to pursue label-free imaging (e.g., brightfield microscopy) as a simple, generic approach for assessing biofilm dynamics.^15,16^ Label-free microscopy has the advantage of being minimally invasive, does not require genetic manipulation to introduce probes, and can potentially be used to assess the visible phenotypes of any bacterial sample, due to the optical contrast intrinsically generated by bacterial cells and biofilm matrix components. Moreover, the simple and affordable optics required for brightfield microscopy are readily available to researchers investigating microbial group behaviors across the world. In this work, we demonstrate that low-magnification brightfield imaging of bacterial culture growth in microtiter plates, together with simple image analysis steps, can be used to reliably quantify microcolony biofilm lifecycles in a sample-agnostic manner. We refer to this approach as “**l**abel-**f**ree **<u>a</u>**nalysis of **<u>b</u>**iofilms” (LFAB). We show that LFAB measurements of biofilm biomass correlate strongly with ground truth confocal microscopy and CV staining measurements across diverse bacterial species. A user-friendly image analysis application is provided, which automates the steps required to extract quantitative data from this assay, so that it can be readily adopted by the field.

As proof of the utility of the LFAB approach, we used high-throughput imaging to discover genes implicated in the biofilm lifecycle of the human pathogen *Streptococcus pneumoniae* (*S. pneumoniae*). *S. pneumoniae* is a major global pathogen, responsible for ∼750,000 annual deaths worldwide, as estimated in 2019.^17^ It forms biofilms on the host epithelium during chronic colonization of the nasopharynx^18–22^ and of the middle ear^23^, as well as during chronic heart disease.^24,25^ These biofilms are sites for strain evolution by horizontal gene transfer^18,26^ and provide *S. pneumoniae* protection from antimicrobials^27,28^ and host immune cells.^18,29,30^ Despite the importance of the biofilm lifecycle for *S. pneumoniae* ecology, few studies have sought to characterize its molecular determinants.^31^ To address this gap, we characterized the microcolony biofilm dynamics of *S. pneumoniae* and identified genes implicated in biofilm development. We found that initial cell inoculum and presence/absence of the cell-surface polysaccharide capsule strongly determined the propensity of an *S. pneumoniae* population to form microcolony biofilms. Moreover, using a transposon screen, we identified novel genes associated with biofilm development. Of these, we validated the role of a locus encoding the cell-surface protein CbpA^32–35^, and the two-component system that regulates it (TCS06)^36–38^, in biofilm formation.^31,39^ In summary, in this study we present LFAB across multiple bacterial species and validate its utility by revealing novel genetic determinants of the *S. pneumoniae* biofilm lifecycle.

## Results

### Low-magnification brightfield microscopy of culture growth yields accurate microcolony biofilm quantifications

In the pathogen *Vibrio cholerae*, individual founding cells develop into discrete biofilm communities at the solid-liquid interface (i.e., the bottom of the well)^10–12^. We refer to these as microcolony biofilms. To assess whether microcolony biofilms could be visualized by brightfield microscopy, we inoculated dilute bacterial cultures (10^4^ cells/mL) into polystyrene-or glass-bottom microtiter dishes and performed time-lapse imaging of static culture growth using a 10x or 4x magnification objective lens (see Methods). Consistent with previous reports, we observed *Vibrio cholerae* microcolony biofilms as areas where light was scattered to a greater degree than their surroundings, yielding darkened regions relative to the background (**Fig. 1A**). We confirmed the verticalization of these communities using confocal microscopy (**Fig. 1B**). To verify that regional darkening in brightfield images is dependent on verticalized biofilm growth, we captured time-lapses of a mutant lacking an exopolysaccharide biosynthesis gene (*vpsL*), known to be required for biofilm formation.^40^ As expected, these cultures exhibited homogenous growth across the field of view with no regional heterogeneity in transmittance (**Fig. 1A**). Notably, we found that other biofilm forming bacteria, including *S. pneumoniae*, *Klebsiella pneumoniae*, and *Pseudomonas* species also exhibited microcolony biofilm formation, with distinct morphologies, at the microplate-liquid interface when grown under similar conditions (**Fig. 1C**).

**Figure 1.**
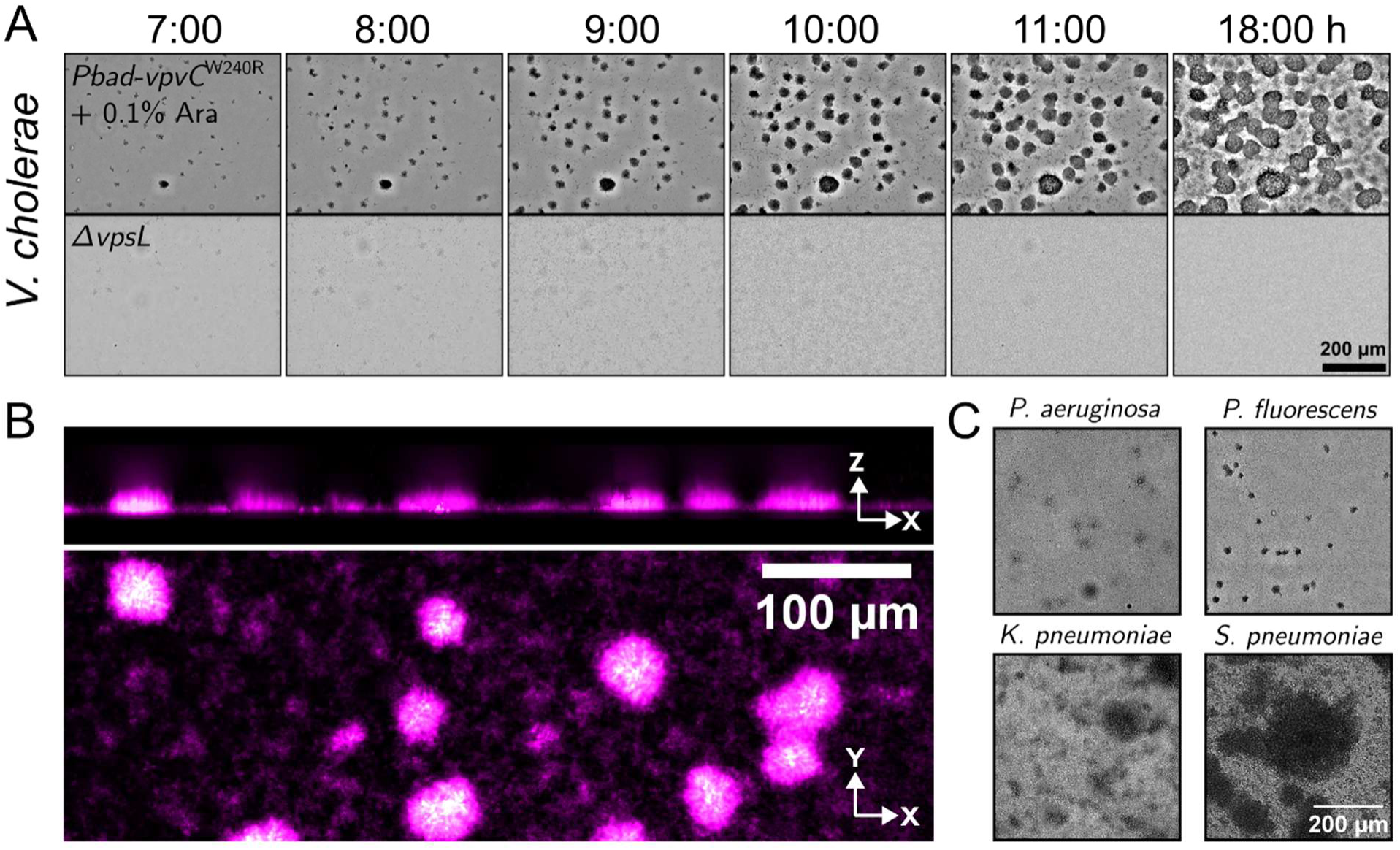
Low-magnification brightfield microscopy captures microcolony biofilm dynamics through regional pixel darkening. **(A)** Time series of brightfield images acquired with a 10x objective for *Vibrio cholerae*. Timepoints reflect the number of hours post-inoculation and scale is as indicated. Top panels represent biofilm formation by a strain carrying *Pbad-vpvC*^W240R^ induced with 0.1% w/v arabinose, which drives expression of a constitutively active diguanylate cyclase, in turn activating biofilm formation^41^. Bottom panels represent the Δ*vpsL* strain, which lacks a *Vibrio* exopolysaccharide gene known to be required for biofilm development. **(B)** Confocal fluorescence images of the induced *Pbad-vpvC*^W240R^ strain after 24 hours of growth, acquired with a 20x magnification objective. Cells were stained with 10µM of the lipophilic dye MM4-64 and images are colored in a magenta-hot lookup table. Top: side-on x-z-projection. Bottom: single x-y plane. Scales and axes are as indicated. **(C)** Brightfield images of the indicated species (strains: *P. aeruginosa* PAO1Δ*wspF, P. fluorescens* SBW25*ΔwspF, K. pneumoniae* KPPR1*ΔwcaJ, S. pneumoniae* SV36 unencapsulated), acquired with a 10x magnification objective at the microplate-liquid interface after 24 hours of culture growth. Scale bar is the same for all species in C and is shown in the bottom right panel.

Our next goal was to exploit the transmittance information present in our low magnification brightfield images to quantify microcolony biofilm biomass, and to compare these measurements to ground-truth biofilm biovolume measured by confocal fluorescence microscopy. We set up an image analysis pipeline towards this goal. As noted above, brightfield images of bacterial cultures exhibited low intensity regions where microcolonies scattered light to a greater degree than the surrounding areas, which were comprised of comparatively sparse planktonic or surface-attached cells (**Fig. 2A**). As a result, we found that microcolony biofilms in our images could be reliably thresholded and their transmittance properties could be measured using simple image analysis techniques (depicted schematically in **Fig. 2A**; for details, see Methods). First, to identify biofilm regions, a local background subtraction was applied to raw images, blurring was performed to eliminate intrinsic intensity variation inside biofilms, and a constant intensity threshold was applied to isolate biofilm regions (depicted as a binary mask on the image). Separately, to convert pixel-wise intensity values in the raw images to optical density (OD) values, a log_10_-based transformation was applied to the transmittance, which accounted for the maximum transmittance (a blank) and the camera noise of the optical configuration. Finally, the binary biofilm mask was applied to the optical density image, yielding a segmented biofilm image, where pixel intensities represent the degree of light scattering within biofilm structures. As a measure of the overall biofilm biomass across the field of view (accounting for biofilm surface area coverage and biofilm transmittance), the average optical density of the segmented biofilm image was computed, which we hereafter refer to as brightfield (BF)-biofilm biomass. We provide a free, user-friendly web application, (see https://github.com/BridgesLabCMU/Brightfield-biofilm-assay), so that our brightfield image analysis pipeline is accessible to all biofilm researchers.

**Figure 2.**
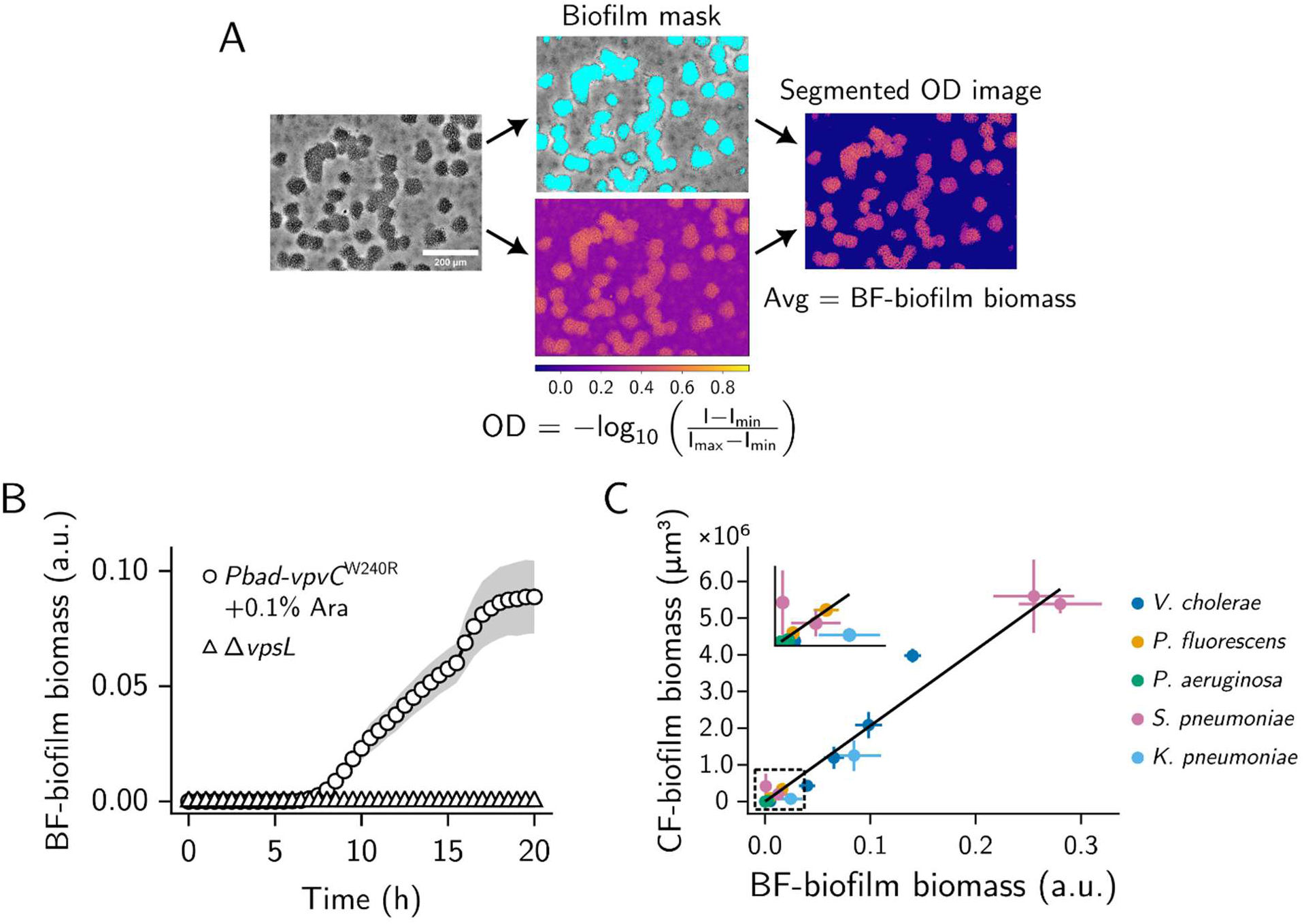
Proof-of-principle for using low-magnification brightfield microscopy of culture growth as a species-agnostic and quantitative biofilm assay. **(A)** Schematic of the pipeline for analyzing brightfield images of culture growth in microtiter plates. Images are from a *V. cholerae* strain carrying a chromosomal *Pbad-vpvC*^W240R^ construct, induced with 0.1% w/v arabinose^41^. Cultures were grown for 24 hours from an initial density of ∼10^4^ cells/mL. Left: raw image. Middle, top: raw image with a binary mask showing biofilm regions overlayed in cyan. The mask was computed as indicated in the main text and Methods. Middle, bottom: optical density image calculated based on the indicated transformation, applied pixel-wise to the raw image. The following represent pixel intensities of respective images: “I” = raw image; “I_min_” = image acquired with the camera shutter closed; “I_max_” = image of fresh media (i.e., a blank). **(B)** Time series of biofilm biomass values, calculated from images of the indicated strains over a 20-hour lifecycle. Lines and scatter points represent means and shading represents standard deviations. **(C)** Correlation of biofilm biomasses, calculated by brightfield microscopy (x-axis) and by spinning-disc confocal fluorescence microscopy (y-axis), for the indicated species (strains: *P. aeruginosa* PAO1 WT andΔ*wspF, P. fluorescens* SBW25 WT andΔ*wspF, K. pneumoniae* KPPR1 WT andΔ*wcaJ, S. pneumoniae* D39 and SV36 encapsulated and unencapsulated, *V. choleraeΔvpsL*, and *Pbad-vpvC*^W240R^ with arabinose concentrations 0%, 0.025%, 0.0375%, 0.05%, and 0.1%). For confocal microscopy, cells were stained with 10 µM (or 20 µM for *S. pneumoniae*) of the lipophilic dye MM4-64. Each point represents the mean biofilm biomass of a strain/arabinose condition. Line represents the best-fit orthogonal distance regression to the data. Error bars represent standard deviation. Inset displays the data points in the boxed region, with re-scaled axes. For brightfield microscopy, *N =* 3 biological with *n =* 3 technical replicates each. For confocal microscopy, *N =* 3 biological replicates. a.u.: arbitrary units. OD: optical density. BF: brightfield. CF: confocal fluorescence. Ara: arabinose. Scale bar as shown.

A major advantage of our approach is that it is non-perturbative, and measurements can be made in real-time throughout biofilm growth. To demonstrate this utility, we captured brightfield images over culture growth and measured biofilm biomass at each timepoint (**Fig. 2B**). Under biofilm inducing conditions in *V. cholerae*, we observed that microcolony biofilm biomass increased steadily post-inoculation, eventually plateauing at ∼18 hours (presumably when nutrients become limited). By contrast, the exopolysaccharide deficient strain exhibited no appreciable microcolony biofilm biomass over the entire growth cycle (**Fig. 2B**). These results suggested that brightfield microscopy of culture growth in microtiter plates, combined with simple image analysis steps, can be used to quantify entire biofilm development cycles in real time.

Having established a pipeline to measure biofilm biomass from brightfield images, we validated our approach by comparing the brightfield quantifications to biofilm biovolume measurements from confocal fluorescence microscopy. Further, we wanted to assess the utility of this pipeline for other biofilm forming bacteria. To this end, we analyzed brightfield images for various biofilm formers under a range of biofilm-inducing conditions (*V. cholerae*, *P. fluorescens*, *P. aeruginosa*, *K. pneumoniae*, and *S. pneumoniae*). Concurrently, we measured the total cellular biofilm biovolume for the same strains/conditions using volumetric spinning-disc confocal fluorescence microscopy, in which biofilm cells were labeled with the lipophilic dye, MM4-64, as a ground-truth for comparison. We found a strong correlation (*r*^2^ = 0.96 based on ordinary least squares) between brightfield and confocal biofilm biomass across all strains and species (**Fig. 2C**). Qualitatively, the correlation between brightfield and confocal biofilm biomass was independent of the objective lens used (as long as sufficiently low magnification objectives lenses were used) (**Fig. S1A**) or microscope (**Fig. S1B**), thus demonstrating that the analyses are agnostic to microscope hardware (see supplemental text and **Fig. S2** for additional discussion of optical considerations for BF-biofilm biomass measurements). Additionally, we found a strong correlation between brightfield biofilm biomass measurements and absorbance due to crystal violet staining (**Fig. S1C**; *r*^2^ = 0.84 based on ordinary least squares). Together, our results show that analyses of biofilm images captured by brightfield microscopy can recapitulate the results of confocal fluorescence microscopy or crystal violet staining and can enable tracking of biofilm lifecycles at high temporal resolution.

### Characterization of *S. pneumoniae* microcolony biofilms using LFAB

To demonstrate the utility of LFAB, we chose to further interrogate the biofilm lifecycle of *S. pneumoniae*. *S. pneumoniae* forms biofilms during chronic colonization of the upper respiratory tract^18–22^, the middle ear^23^, and the heart.^24,25^ Our goal was to resolve the phases of *S. pneumoniae* biofilm development and to identify molecular determinants controlling the lifecycle.

*S. pneumoniae* expresses a polysaccharide capsule on the cell surface, which is a major virulence determinant. Multiple studies have reported an inhibitory role for the capsule in *in vitro* biofilm development.^18,26,28,39,42,43^ *In vivo*, the capsule is downregulated when cells adhere to host epithelial cells, such that low capsule conditions are relevant to colonization^18,44^. Thus, we began by searching for an optimal seeding density in unencapsulated variants of two *S. pneumoniae* strains (model strain D39 and clinical strain SV36). Our aim was to identify conditions that allowed us to assess the ability of individual founder cells to grow into verticalized microcolony biofilms. We captured brightfield time-lapses and observed that when seeding these unencapsulated strains at low density (10^4^-10^1^ cells/mL), distinct cell clusters were observable 5-6 hours post-inoculation and grew radially into microcolonies that plateaued at a peak biomass around 12 hours after inoculation (**Figs. 3A, B and S4A, B**). In contrast, when seeded at high densities (10^5^ cells/mL and above), unencapsulated strains grew into a confluent structure covering the entirety of the field of view. On the other hand, encapsulated strains did not grow into microcolony biofilms at any density, suggesting that their propensity to form verticalized biofilms from single founder cells is reduced (**Fig. S3**). Previous work suggests that encapsulated strains form confluent adherent biofilms at high densities, but we are unable to assess this using the LFAB imaging approach. Finally, while the microcolony phenotype was observed in two phylogenetically distant strains (D39 and SV36), this mode-of-growth may not be universal to *S. pneumoniae*, as the model strain TIGR4 displayed only the confluent phenotype, even when using an unencapsulated strain at low seeding density (**Fig. S5**). We note that the microcolony biofilms examined here are grown under conditions that are quite distinct from the convention in the field. *S. pneumoniae* biofilms have traditionally been examined after inoculation at comparatively high densities using a diverse set of strains, many expressing capsular genes, generally yielding confluent biofilms.^28,42,43,45–47^ These biofilms are often quantified using CV assays^31,46^, or by confocal microscopy.^42,48^ Thus, the *S. pneumoniae* microcolony biofilms analyzed here represent a mode of biofilm growth understudied in this organism. To compare our findings with LFAB to established methods, we performed confocal microscopy and CV assays at low seeding densities with the encapsulated D39 and SV36 strains and their respective unencapsulated mutants. The CV assays and confocal microscopy mirrored the phenotypes observed by LFAB, both for D39 (**Fig. 3C, D**) and for SV36 (**Fig. S4C, D**). We conclude that the *S. pneumoniae* microcolony biofilms captured by LFAB faithfully represent biofilm forming capacity as defined by established assays, while also unlocking new opportunities to study biofilm dynamics in real time and at scale.

**Figure 3.**
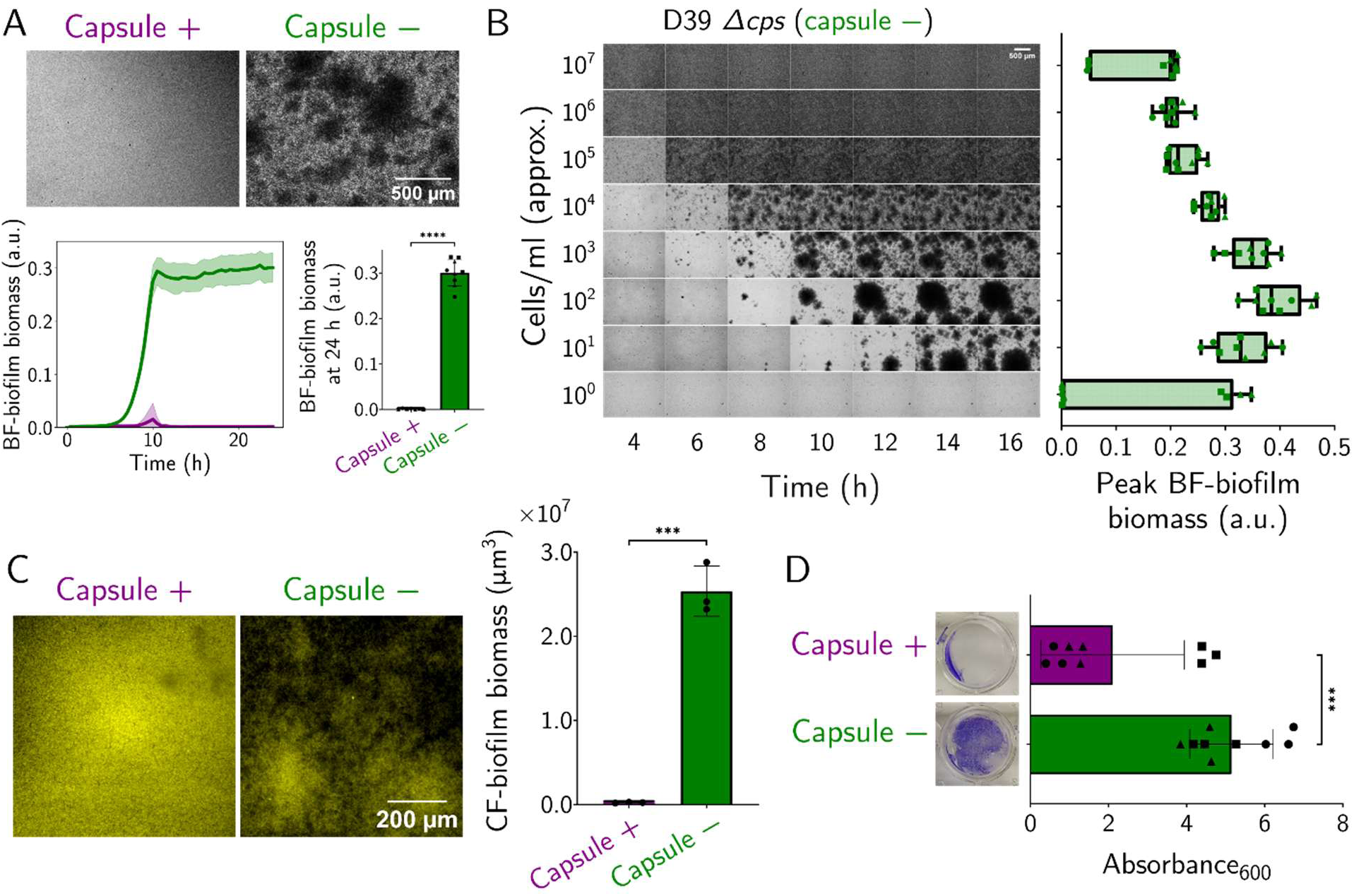
Absence of capsule and low seeding density enable microcolony biofilm formation in *S. pneumoniae* (model strain D39). **(A)** Brightfield microscopy images (4x magnification) of model strain D39 WT (type 2 capsule; left) and isogenic unencapsulated (D39*Δcps*) mutant (right) at 24 hours post seeding, grown from ∼10^3^ cells/mL. Time series plot (bottom) shows microcolony biofilm biomass for WT (purple) and *Δcps* (green) strains, quantified by LFAB from 0-24 hours post seeding at 30 min intervals. Line shows mean and shaded region shows standard deviations. Bar plot (bottom right) shows the 24 hour timepoint. **(B)** Time series brightfield images (left) of D39*Δcps* at various initial cell seeding densities, showing highest seeding density at the top. Points on the boxplot (right) show the peak microcolony biofilm biomass for respective seeding densities across a 24 hour time series. Box plots show median and inter-quartile range, and whiskers show min and max values. **(C)** Confocal microscopy images (showing Z-stack sum projection) of D39 WT (left) and *Δcps* (right) strains. Biofilms were stained with 20 µM of the lipophilic dye MM4-64 and imaged at 24 hours post seeding. Bar plot shows quantified cellular biovolume of microcolonies. **(D)** Crystal violet (CV) assay for D39 WT and *Δcps* at 24 hours post seeding at 20x magnification. Biofilms were washed 3 times with phosphate buffered-saline (PBS), stained with 0.1% crystal violet, and excess stain was removed with 3 more PBS washes. Images show representative wells after staining. CV was then quantified by solubilizing in 70% ethanol and measuring absorbance at 600 nm on the spectrophotometer. For LFAB and crystal violet, *N = 3* biological replicates with *n = 3* technical replicates each; each point on the bar plot is a tech. repl., and each biol. repl. is shown by a unique symbol. For confocal microscopy, *N = 3* biological replicates; each point on the bar plot is a biological replicate. Scale bars are as indicated. (C and D) Bars show mean and error bars show standard deviation. Student’s two-tailed t-test; *** *p < 0.005*. BF: brightfield. a.u.: arbitrary units.

### High-throughput screening for molecular determinants of *S. pneumoniae* microcolony biofilms using LFAB on a transposon library

A major advantage of the LFAB methodology is that it is high-throughput, i.e., it can easily be scaled to image and analyze hundreds to thousands of samples in a day using robotic microscopy. Having established a robust protocol yielding microcolony biofilms that can be reliably quantified using LFAB, we turned towards utilizing the platform’s high-throughput capability to identify genetic determinants of microcolony biofilm formation in *S. pneumoniae*. A previous study reported genes associated with biofilm formation in *S. pneumoniae* using a CV-based screen of a transposon library^31^. Here we revisited this approach using LFAB. To this end, we prepared an indexed library of transposon (Tn) insertion mutants in the unencapsulated model strain D39*Δcps*. We selected this background for two reasons. First, D39 displays high transformation efficiency (>10-fold higher than most clinical strains, including SV36). Second, as a widely used model strain, there is extensive literature in this strain background, allowing for curation of our hits with published data.^49–51^

To screen for biofilm phenotypes, we seeded low-density cultures (diluted to ∼10^3^ cells/mL using a liquid-handling robot) from each of the 2366 Tn-mutant clones in our library, into 96-well microtiter plates. These cultures were grown for 24 hours, after which they were imaged, and biomass was quantified using LFAB. To set the baseline for D39*Δcps* (parental strain of the library) and account for inherent variability between replicates, we imaged and quantified 288 replicates of the parental strain in the same manner (representative images in **Fig. 4B**). We then plotted distributions of biofilm biomass for the parental strain (**Fig. 4A, yellow**) and the transposon clones (**Fig. 4A, green**) in rank order. Using low-stringency thresholds on this distribution, we shortlisted 213 mutant clones that displayed higher or lower microcolony biofilm biomass than the parental strain and performed time-lapse imaging of these shortlisted strains. Based on the time-lapses, we then updated the thresholds to get our final (more stringent) upper and lower significance thresholds (dotted lines in **Fig. 4A**), outside of which biofilm biomass was considered different from the parental strain. This yielded a set of 139 hits (5.9% of the total library size), 69 below and 70 above the parental biofilm biomass threshold. Using Sanger sequencing, we identified the transposon insertion locations in each of these mutants and found that they corresponded to insertions in 123 unique open reading frames. **Table S1** provides a list of all these hits, and **Fig. 4C** displays images of the corresponding microcolony biofilms at 24 hours post-seeding. Visual inspection of the images reveals that the transposon mutants sample the phenotypic space of *S. pneumoniae* microcolony biofilms. They display several distinct microcolony morphologies, with differences in features such as microcolony biofilm size, shape, edge structure, spacing between microcolonies, and distribution of interstitial non-microcolony cells (**Fig. 4C**), which will be quantified in future work. The genes that our screen identified as modulators of *S. pneumoniae* biofilm formation included components with established roles in regulating *S. pneumoniae* biofilms (which validated our approach), as well as genes that have never been characterized (**Table S1**).

**Figure 4.**
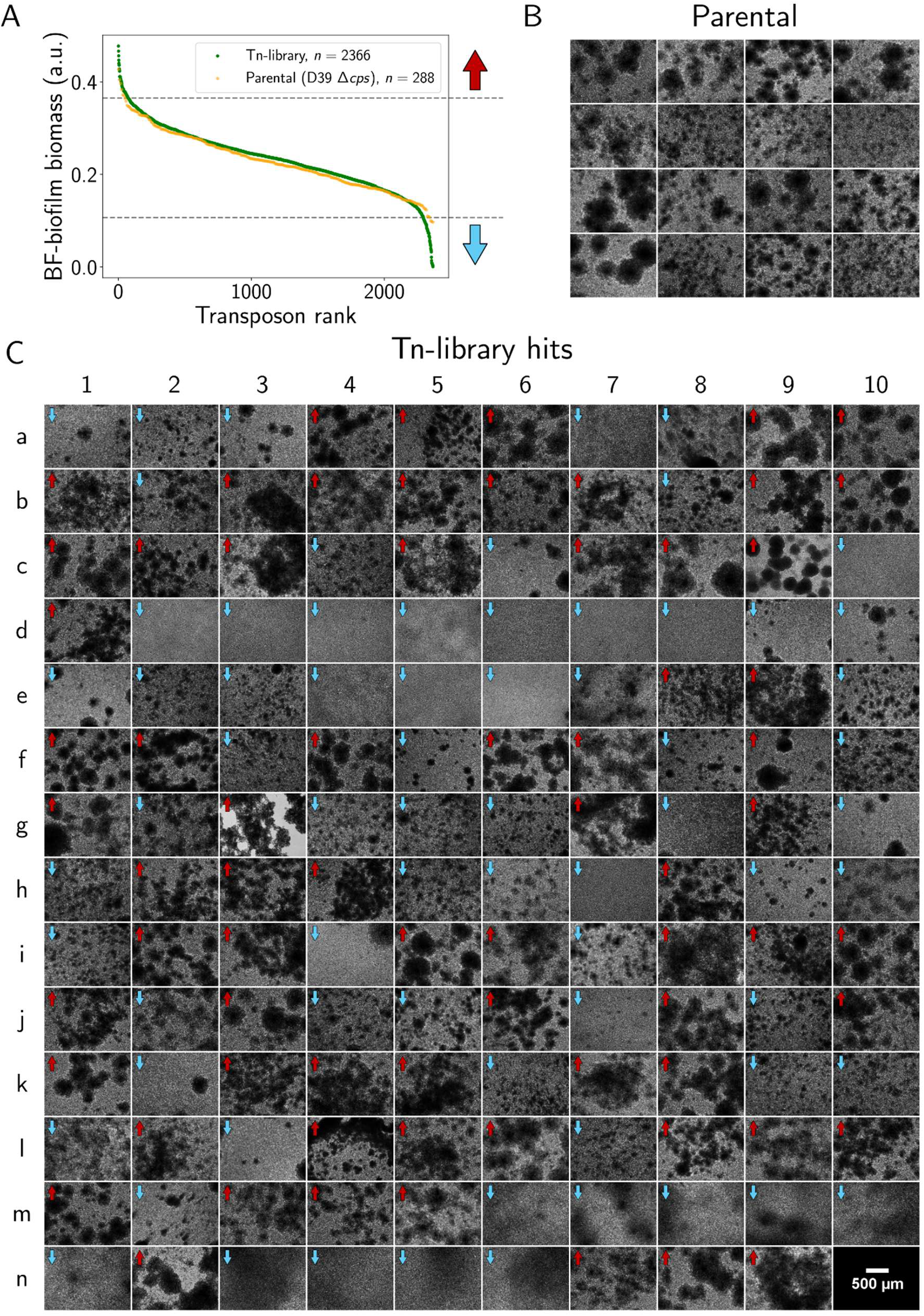
A brightfield microscopy screen identifies novel genes implicated in biofilm development in *S. pneumoniae*. **(A)** Ranked distribution of all Tn-mutant clones screened for microcolony biofilm phenotype at 24 hours post-seeding (green dots). The y-axis displays the biomass quantified by LFAB, and the x-axis shows the rank of each Tn-mutant clone when arranged in decreasing order of biofilm biomass. Yellow points display replicates of the parental strain (D39*Δcps*) imaged in identical fashion to the Tn-library, forming a “null” distribution. Grey dotted lines show the thresholds (0.3651 and 0.1070), on either side of which we considered Tn-mutants to be different from parental strain in microcolony biofilm biomass. **(B)** Representative images (at 24 hours) of parental strain (D39*Δcps*), capturing variability in inoculum due to high-throughput processing by a liquid-handling robot. (**C)** Endpoint (24 hours) images of all 139 shortlisted clones (69 below and 70 above the parental biofilm biomass threshold) from the Tn-library. Details of clones are given in **Table S1**. Blue downward arrow indicates lower microcolony biomass than parental and red upward arrow indicates higher microcolony biofilm biomass than parental. Position n10 is blank and shows the scale bar for all images in panels B and C. BF: brightfield. Tn: transposon.

We identified multiple hits in carbohydrate transporters and modifiers. This class of genes is central to *S. pneumoniae*, which relies exclusively on carbohydrates as a carbon source, a feature reflected in the high number of sugar transporters encoded in its genome.^52,53^ In addition to its critical role as a nutrient, some carbohydrates are also structural components of the capsule^54^ or of the cell wall.^55^ This functional category has been highlighted in previous work^31^, but our screen revealed multiple new components. For example, we found that an insertion in the galactose catabolism gene (*galT*) led to an enhancement in biofilm production (**Fig. 4C position k5**). We also found that an insertion in the cellobiose importer *celB* reduced biofilms (**Fig. 4C position c6**). Most striking was the observation that 7 clones with insertions in a mannitol importer locus displayed a dramatic drop in biofilm mass by 24 hours (**Fig. 4C positions d2-d8**). Overall, the number of hits in carbohydrate-related genes suggests that *S. pneumoniae* links metabolic information to its biofilm lifestyle.

We also observed a dramatic loss of biofilms at 24 hours for insertion within the cell wall biosynthetic enzyme *murM* and its operon (**Fig. 4C positions e5, e6**). This tRNA-dependent aminoacyl transferase plays a critical role in the assembly of branched peptidoglycan on the cell wall.^56,57^ MurM is required for resistance to penicillin^57–59^, highlighting its importance on the cell surface. More recently, MurM has also been shown to buffer entry into the stringent response.^60^ The genes encoding *murM,* as well as the peptidoglycan biosynthesis enzymes *murB* and *murE*, have been previously observed in a screen for biofilm determinants.^31^ Interestingly, we also found that insertion within another cell wall biosynthetic enzyme, the transglycosylase Pbp2a, resulted in increased biofilm biomass (**Fig. 4C position l5**). Together, these findings suggest that genes coordinating cell wall synthesis are linked to biofilm development. Similarly, our screen and a previous screen^31^ both identified cardiolipin synthase, suggesting the composition of the cell membrane may also contribute to biofilm development. For further discussion of hits from our screen, and comparisons to previous results, see Supplemental Discussion. Taken together, applying LFAB screening to *S. pneumoniae* revealed a systems-level connection between fundamental *S. pneumoniae* processes and its biofilm lifecycle.

### Characterization of cell surface protein CbpA and its transcriptional regulators as determinants of microcolony biofilm formation in *S. pneumoniae*

Our final goal was to use LFAB to interrogate a representative pathway identified in our screen. We chose to focus our efforts of characterizing the role of choline binding protein A (*cbpA*) in microcolony biofilm formation because this adhesin is known to play a critical role in *S. pneumoniae* adhesion to host cells, virulence,^32–35^ and has previously been implicated in *S. pneumoniae* biofilms using traditional approaches.^31,39^ Our screen identified 6 mutants with Tn-insertions inside/near *cbpA*, all of which displayed an almost complete loss of microcolony biofilms at 24 hours post-inoculation (**Fig. 4C, positions m6-m10, n1**). In addition, we repeatedly hit genes encoded adjacent to *cbpA*, including a two-component system (*TCS06* or *cbpRS*) and a putative peptide, SPD_2021 (see locus schematic in **Fig. 5A**). To test the role of *cbpA* in microcolony biofilms, we generated a deletion (*ΔcbpA*) and a complement (*ΔcbpA::cbpA^OE^*) in the D39*Δcps* (parental) background. The *ΔcbpA* strain was severely impaired in microcolony biofilm formation (**Fig. 5C**), consistent with the result for the transposon insertions. Analyses of brightfield time-lapses revealed that prior to ∼8-9 h post-inoculation, the *ΔcbpA* and parental strain displayed similar dynamics (**Fig. S6**). However, the *ΔcbpA* strain subsequently deviated from the parental phenotype, resulting in a significantly lower peak biomass value (**Fig. 5C**), and altered biofilm morphology (**Fig. S6**). This phenotype could be complemented by overexpressing *cbpA* from a different genomic location. In fact, the *ΔcbpA::cbpA^OE^* overexpressor strain displayed greater microcolony biofilm biomass than the parental strain and a decrease in interstitial non-microcolony cells (**Fig. 5C**). This phenotype mimicked the parental strain overexpressing *cbpA* (*cbpA^OE^*) (**Fig. 5C**) and demonstrated that elevated expression of *cbpA* gives rise to enhanced microcolony biofilm formation. Together, these data show that CbpA is required for microcolony biofilm formation in *S. pneumoniae*.

**Figure 5:**
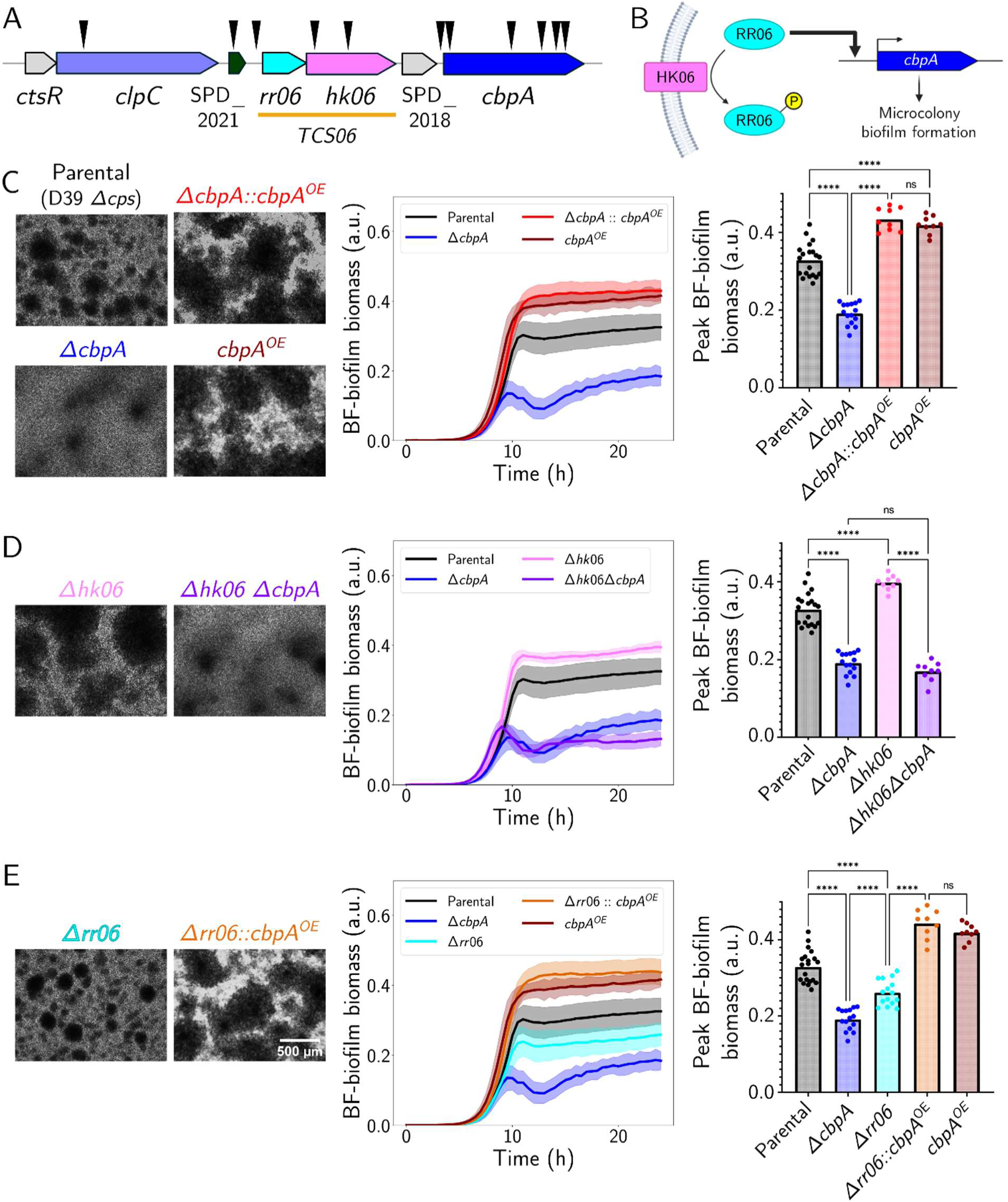
Cell surface protein CbpA and its transcriptional regulation are important determinants of microcolony biofilms in *S. pneumoniae*. (A) Schematic of the locus encoding *cbpA*, *TCS06* and *clpC* in *S. pneumoniae* model strain D39*Δcps*. Black arrowheads show locations of hits from Tn-screen. SPD_2018 & *cbpA* form an operon^37^, and *rr06* & *hk06* form a separate operon^36^, both separate from *ctsR* & *clpC*.^61^ The operon status of SPD_2021 is unknown. **(B)** Schematic showing regulation of *cbpA* expression by the two-component system TCS06. The response regulator (RR06) in unphosphorylated form binds to the *cbpA* promoter region and activates its expression, which in turn promotes microcolony biofilm formation. Once phosphorylated by the histidine kinase (HK06), phosphorylated RR06 can no longer activate *cbpA* expression.^36–38^ (C) Left: Brightfield biofilm images (4x magnification) of the parental strain (D39*Δcps*), *ΔcbpA*, an overexpressor (OE) complement *ΔcbpA::cbpA^OE^* and a parental *cbpA* OE, at 24 hours post seeding. Middle: Time series of microcolony biofilm biomass for the above strains, as quantified by LFAB. Right: Peak microcolony biofilm biomass for each strain. **(D)** As in C, but for *Δhk06* and *Δhk06 ΔcbpA* strains. Data of parental and *ΔcbpA* controls is replotted from A for comparison. **(E)** As in C, but for a *Δrr06* strain and a *cbpA* OE strain in a *Δrr06* background. Data of parental, *ΔcbpA* and *cbpA^OE^* controls is replotted from C for comparison. For all strains, *N = 3-10* biological replicates, with *n = 3* technical replicates each. Scale bars are the same for all images, as shown in D. In time series plots, lines show the mean of all biological and technical replicates, and shaded areas show standard deviations. In bar plots, each data point represents the peak biomass of an individual time series. One way ANOVA, *p < 0.0001* for each set. Šídák’s multiple comparisons test; **** *p < 0.0001*; ns: not significant. BF: brightfield. a.u.: arbitrary units.

Next, we turned our focus to the two-component system encoded adjacent to *cbpA*. TCS06 has been previously shown to transcriptionally regulate *cbpA*.^36–38^ Briefly, this work demonstrated that phosphorylation of the response regulator (RR06) by its cognate histidine kinase (HK06) leads to repression of *cbpA* expression. By contrast, dephosphorylation of RR06 leads to *cbpA* activation (**Fig. 5B**).^36–38^ Therefore, knocking out *hk06* is expected to increase *cbpA* expression (as RR06 is dephosphorylated in this background), whereas knocking out *rr06* is expected to decrease *cbpA* expression.^36^ Consistent with this notion, in our screen a Tn-insertion mapping proximal to the response regulator *rr06* displayed a complete abrogation of microcolony biofilms at 24 hours (**Fig. 4C, position n4**), whereas a clone with insertion inside the histidine kinase *hk06* displayed increased microcolony biofilms and lower background cell density (**Fig. 4C, position n2**). To validate the role of the histidine kinase (*hk06*) and response regulator (*rr06*) in microcolony biofilm formation, we generated clean deletions within this operon in the parental D39*Δcps* background. The phenotypes of these mutants were all consistent with the existing model of *cbpA* regulation. We found that deletion of *hk06*, which increases *cbpA* expression, led to greater microcolony biofilm biomass than the parental strain (**Fig. 5D**). This increase in biomass was dependent on *cbpA* expression, as knocking out *cbpA* in the *Δhk06* background decreased biomass to the same level as the *ΔcbpA* single deletion (**Fig. 5D**). In contrast, we found that deletion of *rr06*, which decreases *cbpA* expression, led to reduced biofilm biomass, displaying an intermediate phenotype between the parental strain and *ΔcbpA* (**Fig. 5E**). These results suggest that the absence of *rr06* does not completely abrogate *cbpA* gene expression. Moreover, overexpression of *cbpA* in the *Δrr06* background rescued the reduced microcolony biomass phenotype, suggesting that the decrease in microcolony biomass in the *Δrr06* strain is due to a decrease in *cbpA* expression (**Fig. 5E**). Further, deletion of both the response regulator and the histidine kinase (*ΔTCS06*), which does not influence *cbpA* levels, did not significantly change the biofilm phenotype (**Fig. S7A, B, C**). Finally, we tested two more hits in this genomic region, a small putative peptide (SPD_2021) encoded upstream of *rr06* and the *clpC* protease, using directed mutations. These phenocopied the wild-type strains, suggesting polar effects of Tn-insertion could be responsible for the observed defects in biofilm development (**Fig. S7A, B, C**). Overall, our studies in the genes surrounding *cbpA* revealed that both *hk06* and *rr06* affect microcolony biofilm formation via their control of *cbpA* expression levels.

In summary, we found that the absence of *cbpA* led to defects in microcolony biofilm formation as well as early disruption of these microcolonies, while high levels of *cbpA* increased microcolony biofilm size and definition. Further, changes in the levels of phosphorylated RR06, which modulates levels of *cbpA,* also drove differences in microcolony biofilm phenotypes. Taken together, using LFAB, we established that CbpA levels are directly associated with microcolony morphology and can be modulated by the individual components of the upstream two-component system TCS06.

## Discussion

Given the limitations of existing methods, here, we asked whether label-free imaging could provide a broadly accessible means to quantify biofilm developmental dynamics. We developed LFAB, which uses low-magnification brightfield microscopy to exploit the intrinsic optical contrast of microcolony biofilms without the need for probes, stains, or genetic manipulation. In LFAB, bacterial populations are initiated at low seeding densities such that biofilm development from founder cells (or small, well-separated groups of cells) can be tracked before complete surface coverage occurs. Because confluence is not reached when microcolonies grow from low seeding densities, the observed biofilm outcomes reflect intrinsic founder-cell biofilm forming capacity rather than homogeneous cell packing in dense populations. Further, since brightfield microscopes are widely accessible to researchers across the world, LFAB can be readily implemented across laboratories and microbial systems. Notably, LFAB occupies a methodological space between crystal violet staining and confocal microscopy: it combines the scalability and affordability of bulk assays with the ability to capture morphological features throughout biofilm development. Strong correlations of LFAB measurements with both crystal violet and confocal benchmarks across diverse species underscores its robustness, and the accompanying automated analysis pipeline makes the approach accessible for broad adoption in studies of microbial community behavior.

As a demonstration of the utility of LFAB, and to further characterize biofilm formation in a notorious pathogen, we applied LFAB to *S. pneumoniae*. We found that, consistent with previous studies, biofilm formation is negatively correlated with capsule production in this organism.^18,26,28,39,42,43^ We then performed a high-content imaging screen which identified 123 candidate genes impacting the biofilm lifecycle of *S. pneumoniae*. The predicted functions for these genes include the transport and metabolism of carbohydrates, the synthesis and autolysis of the cell wall, and cell adhesion, in particular including the cell surface protein CbpA^32–35^ and its regulatory cascade.^36–38^ We show that while CbpA is not required in the early stages of microcolony biofilms, it plays a critical role in the maturation of these communities. CbpA is the most abundant surface protein in *S. pneumoniae*^32^ and is well characterized as an adhesin to host cells.^32–35^ The mechanisms by which CbpA influences biofilms in the absence of host factors remain to be defined. We hypothesize that it facilitates bacterial cell-cell adhesion. Like many surface proteins, CbpA displays a wide range of allelic variants and future studies could test whether and how these variations influence cell-cell contacts, and consequently biofilm maturation and structure.^62,63^ Overall, we present a list of candidate genes involved in *S. pneumoniae* microcolony biofilms, validate the role of CbpA, and demonstrate that LFAB serves as an effective tool to characterize *S. pneumoniae* biofilm determinants.

A previous transposon screen for *S. pneumoniae* biofilm determinants, conducted using the crystal violet assay by the Camilli lab^31^, offers an opportunity to compare our findings to those obtained using traditional biofilm assays. The original study screened both encapsulated and unencapsulated strains seeded at high densities in the TIGR4 strain background (for which we did not measure appreciable microcolony biofilm formation). In the encapsulated strain, aside from capsule biosynthesis genes, this study only identified *lytC* as potentiating biofilms^31^, which we also identified in our screen. In the unencapsulated background, we identified 11 genes common to both sets: choline binding protein A (CbpA)^32–35^, its neighboring histidine kinase^36–38^ and Clp protease^61^, another choline binding protein CbpF^64^, a cardiolipin synthase, ribonuclease Y^65^, an ATP-binding cassette (ABC) transporter, an aldose epimerase, the cell wall synthesis protein MurM^56–59^, and two hypothetical proteins. In addition, we observed a mild biofilm reduction for a *nanA* insertion and the Camilli lab’s study^31^ yielded *nanB*^66^ as a hit. Together, we take the consistency between these screens as proof of principle for the LFAB approach. Our screen also identified 112 unique hits. These include the cell wall peptidoglycan synthesis enzyme MurN^56^, the mannitol transporter system, 3 surface-localized proteins (PavB^67^, NanA^68^ and Pbp2a^69^), the surface charge modifying protein DltA^70^, multiple enzymes involved in carbohydrate processing^52,53^ and general metabolism (e.g., CelB, GalT, GalE, GlgA, PdxS, NrdH, etc.), several carbohydrate transporters in the ABC and phosphotransferase system (PTS) families^52^, a bacteriocin regulator (BlpC)^71,72^ and an immunity protein, an Rgg/SHP cell-cell communication system Rgg1518^73^, the endonuclease EndA involved in DNA uptake^74^, the DNA repair protein RadA^75^, and the cysteine synthase CysK^76^, and numerous proteins with uncharacterized functions.

In summary, studies on *S. pneumoniae* biofilms have revealed that capsule, cell-cell communication, fratricide molecules, nutrient transporters, cell wall synthesis, and adhesins form the genetic toolkit that drives *S. pneumoniae* biofilms, and likely chronic colonization. Yet, a clear playbook of when and how these components are combined to drive formation, maturation, and dispersal of *S. pneumoniae* remain to be determined. Further, it is unclear whether there is one major pathway driving these processes, or whether multiple different pathways achieve similar outcomes. Finally, questions remain as to whether microcolony biofilms or high-density confluent biofilms most closely model the biofilms associated with *S. pneumoniae* carriage, chronic middle ear infections, and heart disease. We propose that LFAB provides a valuable platform to continue studies on the molecular determinants and the pathways that drive biofilms across multiple human pathogens that employ biofilms for colonization and/or infection of the host.

## Methods

### Bacterial growth

All strains used in this work are reported in **Table S2**. *V. cholerae* and *P. fluorescens* strains were propagated on lysogeny broth (LB) plates supplemented with 1.5% agar or in liquid LB with shaking at 30°C. *P. aeruginosa*, and *K. pneumoniae* strains were propagated on LB plates or in liquid LB with shaking at 37°C. *S. pneumoniae* strains were propagated on tryptic soy agar (TSA)-II plates supplemented with 5% sheep blood (BD, BBL, New Jersey, USA; Ref. no. 221261) for overnight growth at 37°C in 5% CO_2_, or in Columbia broth (CB; Remel Inc.; Ref. no. R452972) statically at 37°C in 5% CO_2_.

For LFAB and confocal fluorescence measurements *V. cholerae* strains were grown in M9 minimal medium containing dextrose and casamino acids (1× M9 salts, 100 µM CaCl_2_, 2 mM MgSO_4_, 0.5% w/v dextrose, 0.5% w/v casamino acids), *P. fluorescens* strains were grown in M63 minimal medium containing dextrose and casamino acids (1× M63 salts, 2 mM MgSO_4_, 0.5% w/v dextrose, 0.5% w/v casamino acids), *P. aeruginosa* strains were grown in M63 minimal medium containing arginine (1× M63 salts, 1 mM MgSO_4_, 0.4% w/v L-arginine), *K. pneumoniae* strains were grown in M9 minimal medium containing dextrose (1× M9 salts, 100 µM CaCl_2_, 1 mM MgSO_4_, 0.4% w/v dextrose), and *S. pneumoniae* strains were grown in CB. For *K. pneumoniae*, the M9 medium containing dextrose (but no additional salts) was chelated for 3 hours using Chelex 100 chelating resin (BioRad catalog no. 1421253). After filter sterilization, salts were added.

### Biofilm growth

For *V. cholerae*, *P. fluorescens*, and *P. aeruginosa*, unless otherwise indicated, strains were grown overnight in LB and subsequently back-diluted to ∼10^4^ cells/mL in minimal medium.

For *K. pneumoniae*, overnight LB cultures were washed 1x in PBS, then back-diluted to ∼10^4^ cells/mL in minimal medium. For *S. pneumoniae* strains, cultures were grown overnight on TSA-II plates with 5% sheep blood, at 37°C in 5% CO_2._ In the morning, cultures were inoculated into CB at OD_600_ ∼0.02-0.03, and grown to OD_600_ = 0.05, i.e. early log phase (corresponding to ∼10^7^ cells/mL). Cultures were then back-diluted in fresh CB to ∼10^3^ cells/mL unless otherwise indicated. 200 µL of diluted cultures were subsequently grown statically in 96-well microtiter plates (Costar; Ref. no. 3370). For experiments pertaining to Fig. 1, *V. cholerae, P. fluorescens,* and *P. aeruginosa* biofilms were grown for 24 hours and subsequently washed 6x in PBS media prior to imaging. The procedure was identical for *K. pneumoniae* but without wash steps to avoid biofilm disruption. For *S. pneumoniae*, the images were acquired at 10 h post seeding without washing, to capture the peak stage of microcolony biofilm growth.

### Microscopy

Brightfield time-lapse microscopy images were acquired at 30-min intervals at 30 °C (*V. cholerae*, *P. fluorescens*, *P. aeruginosa*, and *K. pneumoniae*) or 37 °C (*S. pneumoniae*) on an Agilent Biotek Cytation 1 imaging plate reader using either a 10x air objective (Olympus Plan Fluorite, NA 0.3) or a 4x air objective (Olympus Plan Fluorite, NA 0.13) driven by Biotek Gen5 (Version 3.12) software. In Fig. S1, images were additionally acquired on an EVOS M5000 imaging system (Invitrogen, Thermo Fisher Scientific Inc.), using an EVOS Plan phase-contrast 10x air objective (0.30NA/7.13WD), or using a Nikon Ti-E inverted epifluorescence HCS imager, equipped with a Photometrics Evolve EMCCD camera and a 10x air objective, as indicated. Samples were grown in 96-well polystyrene microtiter plates (Costar; Ref. no. 3370) or in Mattek glass bottom 96-well microtiter plates (MatTek Corporation; Part no. P96G-1.5-5-F). Spinning disc confocal images were acquired using a motorized Nikon Ti-2E stand outfitted with a CREST X-Light V3 spinning disk unit, a back-thinned sCMOS camera (Hamamatsu Orca Fusion BT) and a 20x air objective (Nikon Plan Apochromat, NA 0.75) driven by Nikon Elements software (Version 5.42.02). The source of illumination was an LDI-7 Laser Diode Illuminator (89-North). Cells were stained with 10 µM (or 20 µM for *S. pneumoniae*) of the lipophilic dye MM4-64 (AAT Bioquest) for 1 hour and were imaged with an excitation wavelength of 561 nm. *V. cholerae, P. fluorescens*, *P. aeruginosa*, and *K. pneumoniae* strains were grown in 18-well glass-bottom chamber slides (ibidi GmbH). *S. pneumoniae* strains were grown in a Mattek glass-bottom 96-well microtiter plate (MatTek Corporation; Part no. P96G-1.5-5-F).

### LFAB image analysis and web application

All code for image analysis were written in the Julia programming language (v1.11.2) and are publicly available on GitHub (https://github.com/BridgesLabCMU/Brightfield-biofilm-assay). All brightfield images were subjected to the same image analysis pipeline. A web application interface for performing image analysis can be used, and the instructions are provided at the GitHub page.

To generate a mask (i.e., a spatial indicator) of biofilms in a brightfield image or timeseries of images, a corresponding “background” image is generated through the application of a large-kernel Gaussian filter (default side length = 100 pixels in *x* and *y*; for *S. pneumoniae* and *K. pneumoniae*, we used side length = 200 pixels in *x* and *y* due to large biofilm sizes). The background image is then subtracted from the raw image, normalizing the local contrast within an image and, for time-lapse imaging, normalizing the background intensity across timepoints. At this point, time-lapses are registered (aligned) using the single-step discrete Fourier transform algorithm described by Guizar-Sicairos and colleagues to correct for frame-to-frame jitter.^77^ A small-kernel Gaussian filter (side length = 2 pixels in *x* and *y*) is then applied to control for intensity variation within biofilms, and a constant threshold is applied to generate a segmented mask of biofilms. In our study, the following thresholds were used for respective organisms: 0.03 (*S. pneumoniae* and *K. pneumoniae*) and 0.04 for all other species. If dust correction is selected (time-lapses only), all pixels in the mask that are 1’s at the beginning of the time-lapse and transition to 0’s at any point are taken to be dust and are set to 0 throughout. If closed-shutter and blank images (I_min_ and I_max_, respectively) are not supplied, the mask is directly applied, pixel-wise, to the raw image, and the average value in the resulting image yields the “biofilm biomass” value. All *S. pneumoniae* data was analyzed in this manner, without supplying I_min_ and I_max_ images. If I_min_ and I_max_ images (or just I_min_ if the first image of a time-lapse is taken to be a blank image) are supplied, an optical density image is calculated as

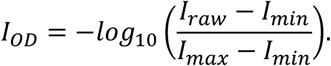

The mask is then applied, pixel-wise, to the optical density image, and the average value for all pixels in the resulting image yields a “biofilm biomass” value. For display purposes (in particular, for time-lapses, where the background intensity might change dramatically due to growth of planktonic cells), local contrast and global (temporal) background intensity are normalized using a local averaging filter based on integral arrays.

### Confocal fluorescence microscopy image analysis

Confocal fluorescence images were acquired as described above. For each image, we reduced the contribution of Poisson shot noise in the images using the Denoise.ai module available in Nikon Elements software (Version 5.42.02). We then performed top-hat filtering of each x-y slice to reduce the contribution of out-of-focus light along the optical axis. A z-projection was computed and then blurred using a Gaussian filter (side length = 10 pixels). A bounded automatic threshold was then applied to the z-projection using Otsu’s method, generating a 2D mask. In effect, this procedure isolated regions in the *x*-*y* plane that contained biofilms (regions of locally greater thickness than the surroundings). Then, for each slice in the original tiled z-stack (corrected for anisotropy along the optical axis), a mask of cells was computed using a bounded automatic threshold, again using Otsu’s method. The z-projection mask was then applied to the cell mask, and the sum of the pixels in the resulting binary image yielded a value for the cell area occupied by biofilms. The sum of these areas over all slices yielded a total biofilm biomass for the 3D image.

### Crystal violet assay

For *S. pneumoniae*, biofilms were seeded at 10^3^ cells/mL into 12-well microtiter plates (VWR USA; catalog no. 10062-894) using CB as growth media, and grown for 24 h. At endpoint, supernatant was removed and biofilms were gently washed 3 times with phosphate buffered-saline (PBS). Biofilms were then incubated with 0.1% crystal violet (CV) solution for 10 min at room temperature. Excess stain was removed with 3 more PBS washes. CV was then quantified by solubilizing in 70% ethanol and measuring absorbance at 600 nm on the spectrophotometer. The protocol for *V. cholerae* was identical except that cultures were grown in M9, as described above, in 24-well plates. Readings were taken at 590 nm on the spectrophotometer.

### Bacterial construct generation and transformation

For generating *S. pneumoniae* mutants, we used a two-step transformation approach to generate clean deletions using the **f**luorescent **<u>r</u>**emovable **<u>an</u>**tibiotic **<u>c</u>**assette (FRANC) containing spectinomycin-resistance *aad9*, sucrose-sensitivity *sacB* and a fluorescent mCardinal marker.^78^ Briefly, for the first transformation FRANC was inserted to replace the region desired for deletion. Colonies were screened for spectinomycin resistance and mCardinal fluorescence. In the second step, the FRANC was removed using a construct with only the two flanking regions. Colonies were selected for sucrose resistance.

In brief, around 2 kb of flanking regions up-and downstream of the gene/region of interest, and FRANC were amplified by PCR using Q5 2x Master Mix (NEB, USA). PCR fragments were verified by gel electrophoresis and subsequently cleaned up via DNA Clean & Concentrator-5 kit (Zymo Research, USA). Fragment assembly was carried out using an in-house GIBSON assembly mix. All primers used for construct generation are listed in **Table S3**.

Strains were inoculated in CB and grown to an OD_600_ of 0.05. Then 1 ml culture was incubated with the previously assembled DNA construct and 125 ng CSP-1 peptide (EMRLSKFFRDFILQRKK, GenScript, USA). After a 2 hours incubation period, cultures were plated on respective CB agar spectinomycin plates (100 µg/mL spectinomycin) or CB agar sucrose plates (10% w/v sucrose). After overnight incubation, resistant colonies were picked, regrown in CB media with 100 µg/mL spectinomycin or 10% sucrose (w/v) and frozen stocks were generated. Mutants were verified via PCR and Sanger sequencing.

### Isolation/generation of unencapsulated *S. pneumoniae* strains

The unencapsulated version of the SV36 strain was isolated as a spontaneous mutant in the lab. SV36, which expresses the type 3 capsule, makes visibly thick mucoidal colonies on TSA+blood plates. Spontaneous mutants that do not express the capsule make visibly distinct non-mucoidal colonies. One such non-mucoidal colony was picked, propagated and its genome was sequenced. We found a single nonsense mutation in the *cpsA* gene. This strain was used as the unencapsulated SV36 in this study.

The unencapsulated D39 was engineered by making a markerless gene deletion using the FRANC-based process described above. The deletion spanned multiple capsule genes (*cpsA-H* and *cpsT*) and mirrored the natural deletion found in the unencapsulated lab strain R6. This engineered strain was used as the unencapsulated D39 in this study.

### Transposon library generation and screening

The indexed transposon (Tn) library was prepared using genomic DNA from a pre-existing pooled Tn-mutant library in the D39 WT (encapsulated) background, by transforming it into D39*Δcps* and picking single colonies off a selection plate.^79^ Each colony was picked into an individual well of a 96-well plate containing Columbia Broth (CB), grown for 8 h at 37°C in 5% CO_2_, and frozen stocks were generated. The library consisted of 25 such 96-well plates.

To grow the library subsequently for screening, an autoclaved PCR plate was used as a “stamp” to inoculate from each frozen stock plate into a fresh 96-well plate containing CB. These were grown for 8 h at 37°C in 5% CO_2_, and a liquid handling robot (Opentrons) was used to dilute down to 10^-5^ for seeding biofilms into another 96-well plate. The biofilms were grown for 24 h, and imaged at endpoint on the Agilent Biotek Cytation 1 imaging plate reader using a 4x magnification objective.

### Identification of transposon insertion location by arbitrary PCR and Sanger sequencing

To identify location of transposon inserts, a PCR reaction was run using Q5 2x Master Mix (NEB, USA), with a forward primer inside the Tn sequence (P1) at 0.5 µM concentration, a random reverse primer mixture containing a defined overhang (adapter) sequence (P2) at 10x concentration of P1 (5 µM), and a cell suspension of the Tn-mutant as template. The following PCR conditions were used: 98°C for 30 s + {98°C for 15 s + 30°C for 30 s + 72°C for 30 s} x 5 cycles + {98°C for 15 s + 55°C for 30 s + 72°C for 30 s} x 30 cycles + 72°C for 2 min. The product of this first PCR was used as a template to run a second PCR (conditions: 98°C for 30 s + {98°C for 15 s + 55°C for 30 s + 72°C for 30 s} x 30 cycles + 72°C for 2 min), using another forward primer (P3) inside the Tn sequence (downstream of where P1 binds), and a reverse primer that binds to the overhang adapter sequence in P2. Following the second PCR, the amplified fragment was purified by DNA Clean & Concentrator-5 kit (Zymo Research, USA). It was Sanger sequenced using primer P3, by Genewiz from Azenta Life Sciences, USA.

Upon receipt of the Sanger sequencing read, the part of the read that matched the end of the Tn was trimmed off. The trimmed read was used as a query to run nucleotide BLAST against the *S. pneumoniae* D39 genome on NCBI (RefSeq accession GCF_000014365.2). The region of best match (start coordinate of BLAST hit) was recorded as the insertion location of the transposon. The locus was inspected visually, the nearest gene in the RefSeq annotation was identified, and the gene IDs (new NCBI ID “SPD_RSxxxxx” and old ID “SPD_xxxx” were recorded. Corresponding R6 IDs “sprxxxx” were identified using PneumoBrowse.^49^

## Supporting information

Table S1: Tn-screen hits

## Acknowledgements

We thank Dr. Carlos Orihuela for sharing TIGR4 (encapsulated and unencapsulated) strains and insightful comments. We thank Dr. Vaughn Cooper for sharing *P. fluorescens* strains, Dr. Laura Mike for *K. pneumoniae* strains, and Dr. Catherine Ambruster for *P. aeruginosa* strains. We thank Dr. Jason Rosch for his contribution to reagents and protocols associated with preparation of the transposon library. Figure 5B was created in BioRender (https://BioRender.com/34glawf).

This work was supported by NIH grant R00AI158939, a Shurl and Kay Curci Foundation grant (https://curcifoundation.org/), a Kaufman Foundation New Investigator Research Grant KA2023-136488 (https://kaufman.pittsburghfoundation.org/), a Damon Runyon Cancer Research Foundation Dale F. Frey Award for Breakthrough Scientists 2302-17 (https://www.damonrunyon.org/), and startup funds from Carnegie Mellon University to AAB. It was also supported by NIH grant R21AI182784 (NLH), a Kaufman Foundation New Initiative research grant KA2024-144002 (NLH), the Carnegie Mellon’s Center for Machine Learning and Health at the School of Computer Science Fellowship (PMR), and support from the department of Biological Sciences at CMU.

## Supplemental Text

During image acquisition, a relevant control parameter is the focal position. We found that when imaging small biofilms, biofilm biomass values were affected considerably by focal position, whereas the effect was attenuated when imaging larger biofilms **(Fig. S2A)**. Thus, establishing a single focal position is necessary to ensure consistency across experiments that would enable quantitative comparisons. Moreover, we found that in cases where populations were comprised of a large fraction of planktonic cells, planktonic cell light scattering led to lower biofilm biomass values due to their homogenizing effect on local image intensities, which impedes detection of microcolony biofilms **(Fig. S2B)**. Thus, we caution against an interpretation of the absolute fraction of dispersed cells in cases where populations undergo biofilm maturation-dispersal lifecycles. In cases where it is desirable to compute such fractions, we recommend endpoint washing. Planktonic cell effects, together with focal position effects, comprise the major sources of non-biological quantitative variance within the LFAB framework.

## Supplemental Discussion

We also identified the autolysins *lytB* and *lytC* in our screen. This is consistent with previous studies, where the murein hydrolases LytA, LytB and LytC have been shown to contribute to biofilms.^31,39,80^ Autolysins may contribute by lowering the number of cells and/or by increasing extracellular DNA (eDNA) within the biofilm.^39^ Of note, in our study, the *lytB* insertion yielded a dramatic phenotype characterized by increased biomass and large asymmetrical colonies, with the vast majority of the cells in field of view being concentrated within the colonies. (**Fig. 4C, position g3)**. LytB promotes cell separation, such that defects in this protein lead to longer chains.^81–83^ We propose that increased chaining in the *lytB* transposon insertion promotes an architecture that resists shear and traps cells and matrix. Alternatively, changes in *lytB* may affect biofilms indirectly by driving hyper-competence^84^, in turn promoting changes in gene expression such as upregulation of BriC, a peptide linked to biofilm development.^85^ Akin to autolysins, fratricidal molecules also contribute to cell death. Our screen revealed decreased biofilm biomass by Tn-insertion in *blpC* **(Fig. 4C position e1)**, the quorum sensing pheromone that drives bacteriocin production.^71,72^ While cells in our study are clonal, and thus devoid of inter-strain competition, a study by Aggarwal and colleagues demonstrated that BlpC plays a role in colonization in a clonal population. Cells that stochastically express *blpC* early, target identical cells that have not yet expressed bacteriocin(s) and their associated immunity protein.^86^ It is plausible that similar dynamics are also at play in our microcolony biofilms, where BlpC could drive intra-strain lysis and accumulation of eDNA early in microcolony biofilm development.

Beyond the autolysins, another category of hits in our screen was cell surface molecules. Neuramidase A (*nanA*) is a sialidase^68^, which contributes to colonization and biofilm development by cleaving sialic acid residues from host cells.^45,87^ It also encodes a lectin-like domain that affords it additional adhesive properties.^88^ We found that *nanA::Tn* exhibited decreased microcolony biofilm biomass **(Fig. 4C position j4)**. Similarly, a previous study on confluent biofilms found that disruption of *nanB*, a gene encoding a second sialidase, also hindered biofilm development.^31^ Among other surface proteins that we identified was the pneumococcal adherence and virulence factor B (*pavB*), a fibronectin binding surface protein^67^, whose insertional mutant showed increased microcolony biofilm biomass (**Fig 4C position a10**). Tn-insertions were also observed in the cell surface adhesin *cbpA*^32–35^ and an adjacently-encoded regulatory two-component system *TCS06*^36–38^, which we have further characterized in the Results section. The presence of host-binding proteins in our screen suggests these cell-surface proteins have additional biofilm-related functions that are independent of host cells.

## Supplemental Figures

**Figure S1.**
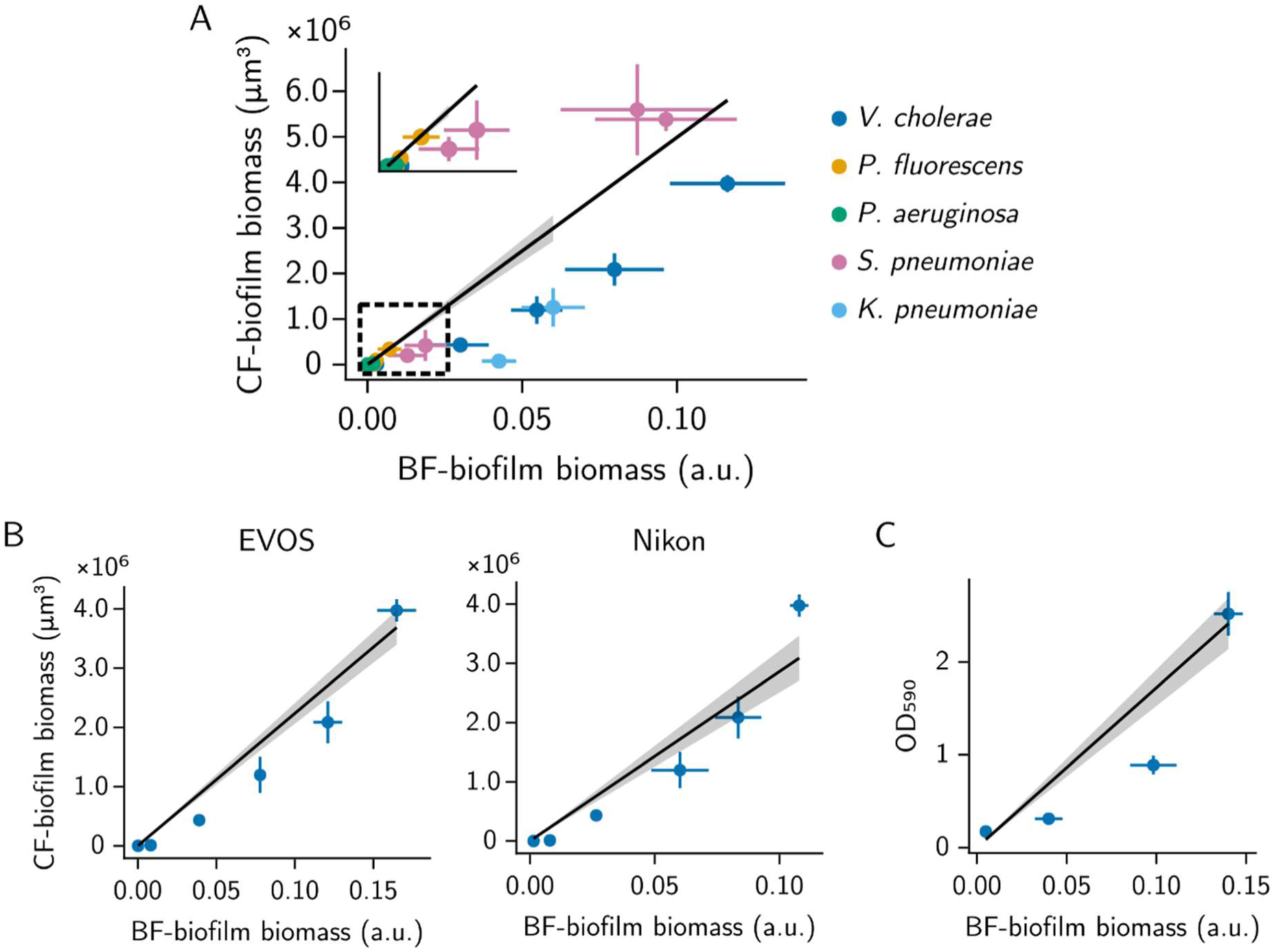
LFAB correlations with confocal microscopy and crystal violet methods across microscopes and objectives. In all panels, measurements were taken after 24 hours of growth. **(A)** Correlation of biofilm biomass calculated by brightfield microscopy (10x objective) on an Agilent Biotek Cytation 1 imaging plate reader and by spinning-disc confocal microscopy for the indicated species. Each point represents a separate strain or biofilm induction condition. Line represents the best-fit orthogonal distance regression to the data. *r*^2^ = 0.77 based on ordinary least squares fit to the data. Inset displays the data points in the boxed region, with re-scaled axes. Error bars represent standard deviation. For brightfield biofilm biomass measurements, *N =* 3 biological and technical replicates. For confocal fluorescence biofilm biomass measurements, *N =* 3 biological replicates. **(B)** As in A but where brightfield biofilm biomass was measured on an EVOS (left) or Nikon (right) microscope. *r*^2^ = 0.95 and *r*^2^ = 0.92, respectively, based on ordinary least squares. **(C)** As in A, B but a correlation between brightfield biofilm biomass and crystal violet staining for *V. cholerae (N =* 3 biological replicates). *r*^2^ = 0.84. To achieve variation in biofilm biomass production in *V. cholerae* strain chromosomally encoding a *Pbad-vpvC*^W240R^ construct was used. This construct, when induced with arabinose, drives expression of a constitutively activated diguanylate cyclase, which in turn activates biofilm formation.^41^ The strain was induced with varying concentrations of arabinose. a.u.: arbitrary units. BF: brightfield. CF: confocal fluorescence.

**Figure S2.**
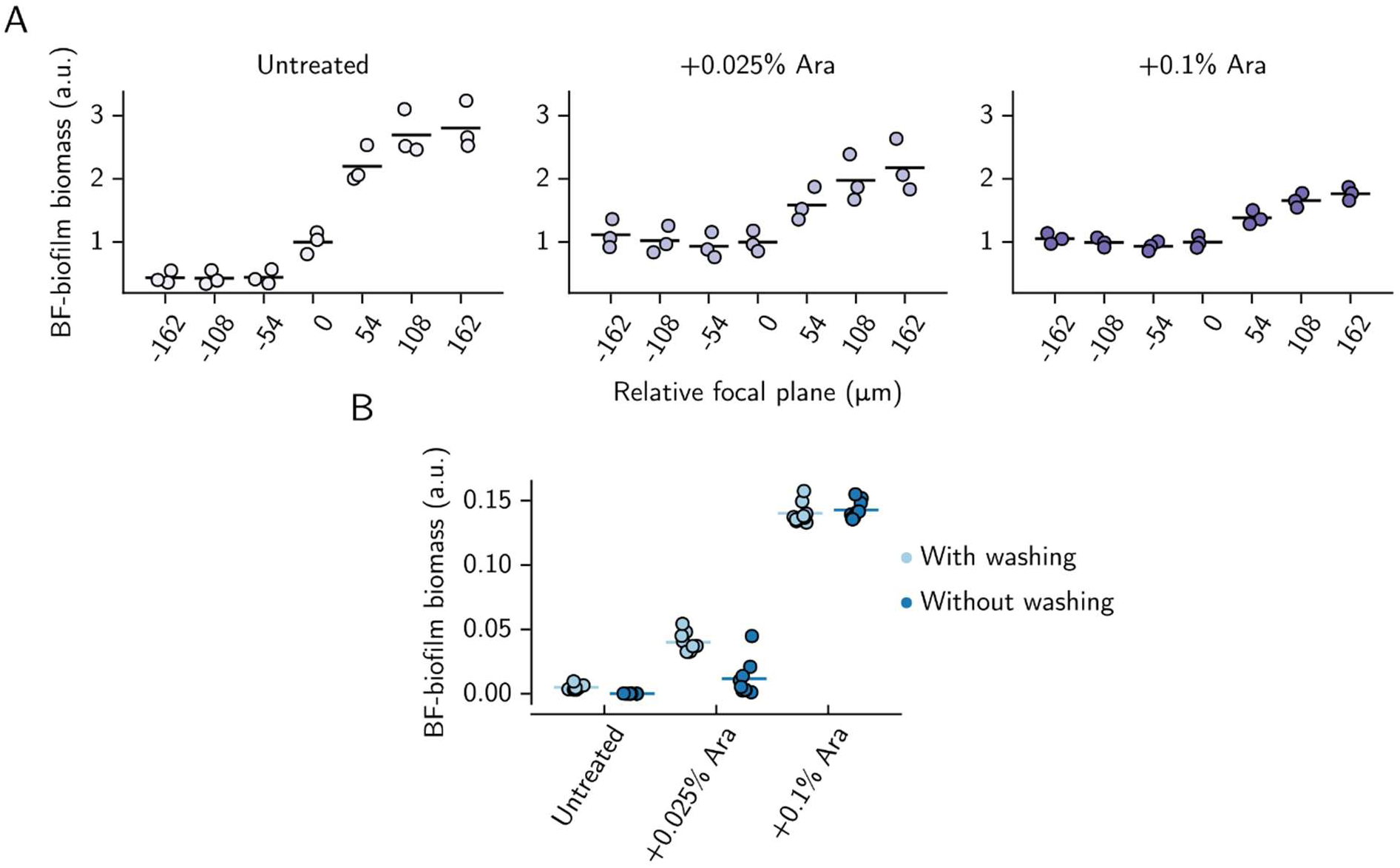
**Technical considerations for the LFAB approach**. **(A)** Left: biofilm biomass of a *V. cholerae* strain carrying a chromosomal *Pbad-vpvC*^W240R^ construct^41^, without induction, as measured from brightfield images at the indicated focal planes. A focal plane of 0 represents an in-focus image. Data are normalized to the mean biofilm biomass of the in-focus images. Each point represents a replicate for *N =* 3 biological replicates. Middle, Right: as in the left panel for the same strain induced with the indicated arabinose concentrations. **(B)** Brightfield biofilm biomass of the same *V. cholerae* strain as in A, grown in the presence of the indicated concentrations of arabinose, and with or without an additional wash step as indicated. Each point represents a replicate for *N =* 3 biological and technical replicates. Increasing levels of *Pbad-vpvC*^W240R^ induction result in increasing levels of biofilm biomass production.^41^ a.u.: arbitrary units. BF: brightfield. Ara: arabinose.

**Figure S3.**
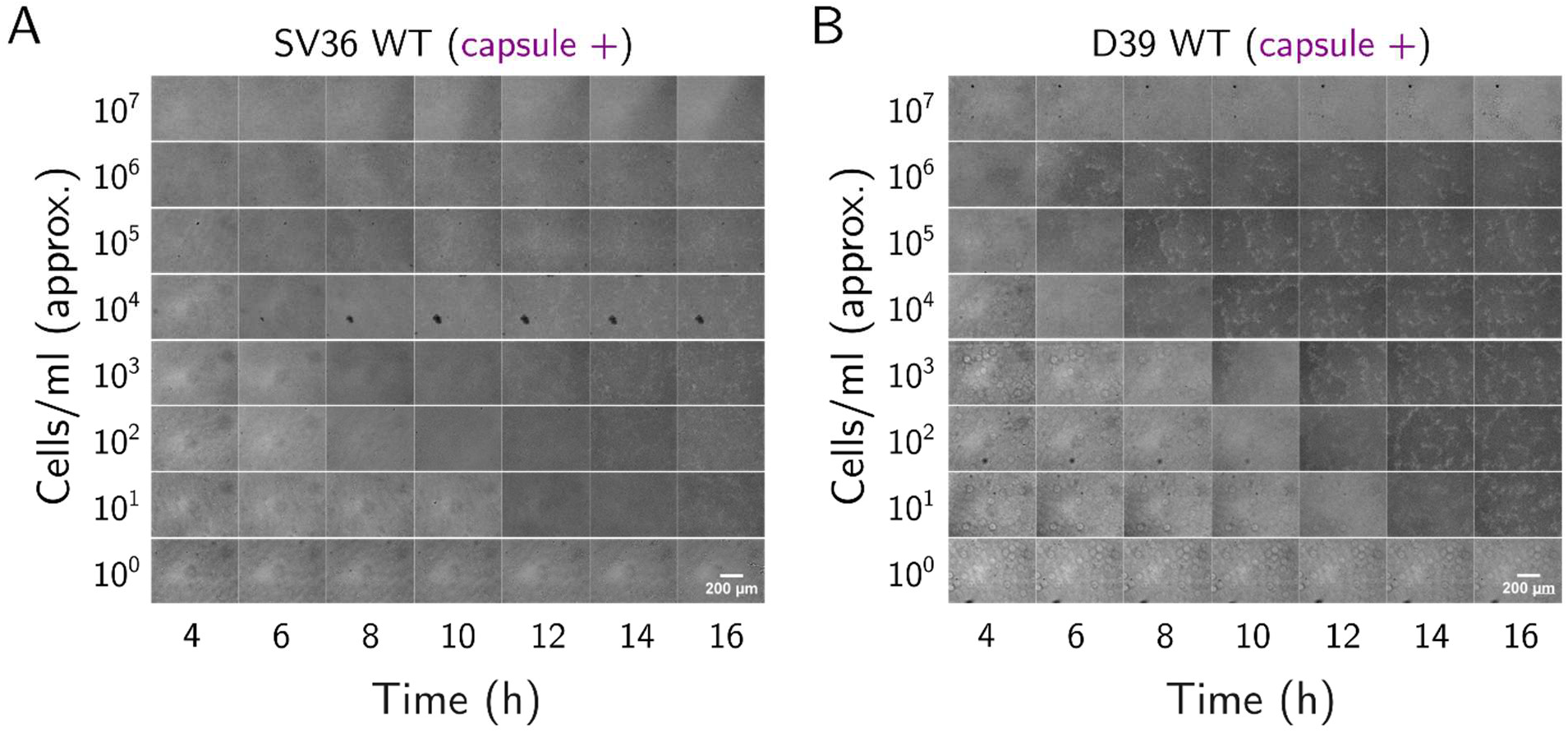
Encapsulated WT strains SV36 and D39 form confluent biofilms at all seeding densities. **(A)** Time series brightfield images of SV36 WT (encapsulated; type 3 capsule) at various initial cell seeding densities, showing highest cell inoculum at the top. **(B)** As in A, for D39 WT (encapsulated; type 2 capsule). All images were acquired at 10x magnification. *N = 3* biological replicates with *n = 3* technical replicates each. Scale bar is the same for all images, and is indicated on the bottom right of each panel.

**Figure S4.**
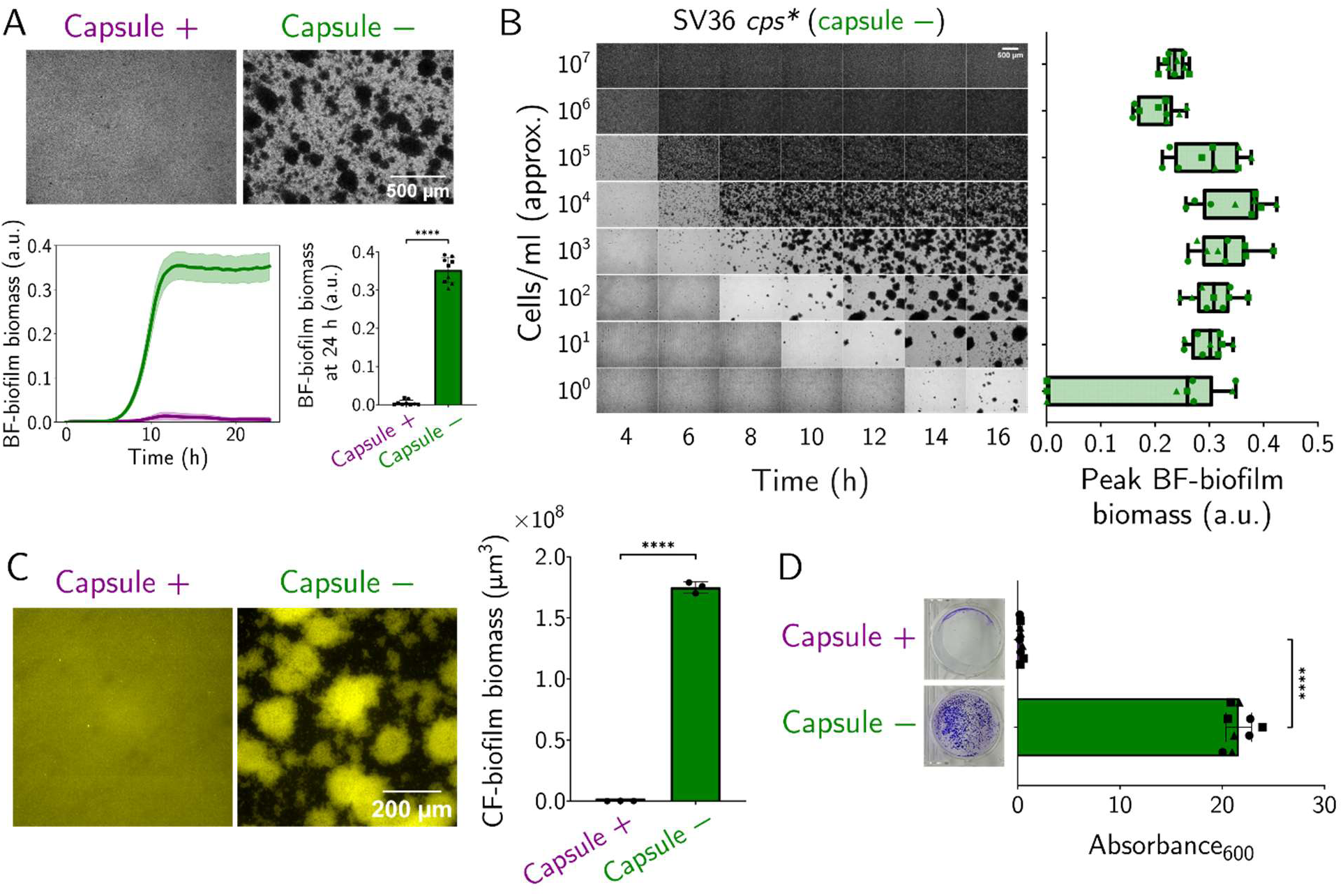
Absence of capsule and low seeding density enable microcolony biofilm formation in *S. pneumoniae* (clinical strain SV36). **(A)** Brightfield microscopy images (4x magnification) of clinical strain SV36 WT (type 3 capsule; left) and isogenic unencapsulated mutant (*cps*\*; right) at 24 hours post seeding, grown from ∼10^3^ cells/mL. Time series plot (bottom left) shows microcolony biofilm biomass for WT (purple) and *cps*\* (green) strains, quantified by LFAB from 0-24 hours post seeding at 30 min intervals. Line shows mean and shaded region shows standard deviations. Bar plot (bottom right) shows the 24 hour timepoint. **(B)** Time series brightfield images (left) of SV36 *cps*\* at various initial seeding densities, showing highest seeding density at the top. Points on the boxplot (right) show the peak microcolony biofilm biomass for respective inocula across a 24 hour time series. Box plots show median and inter-quartile range, and whiskers show min and max values. **(C)** Confocal microscopy images (showing Z-stack sum projection) of WT (left) and *cps*\* (right) SV36 strains. Biofilms were stained with 20 µM of the lipophilic dye MM4-64 and imaged at 24 hours post seeding at 20x magnification. Bar plot shows quantified cellular biovolume of microcolonies. **(D)** Crystal violet (CV) assay for SV36 WT and *cps*\* at 24 hours post seeding. Biofilms were washed 3 times with phosphate buffered-saline (PBS), stained with 0.1% crystal violet, and excess stain was removed with 3 more PBS washes. Images show representative wells after staining. CV was then quantified by solubilizing in 70% ethanol and measuring absorbance at 600 nm on the spectrophotometer. For LFAB and crystal violet, *N = 3* biological replicates with *n = 3* technical replicates each; each point on the bar plot is a tech. repl., and each biol. repl. is shown by a unique symbol. For confocal microscopy, *N = 3* biological replicates; each point on the bar plot is a biol. repl. Scale bars are as indicated. (C and D) Bars show mean and error bars show standard deviation. Student’s two-tailed t-test; **** *p < 0.0001*. BF: brightfield. a.u.: arbitrary units.

**Figure S5.**
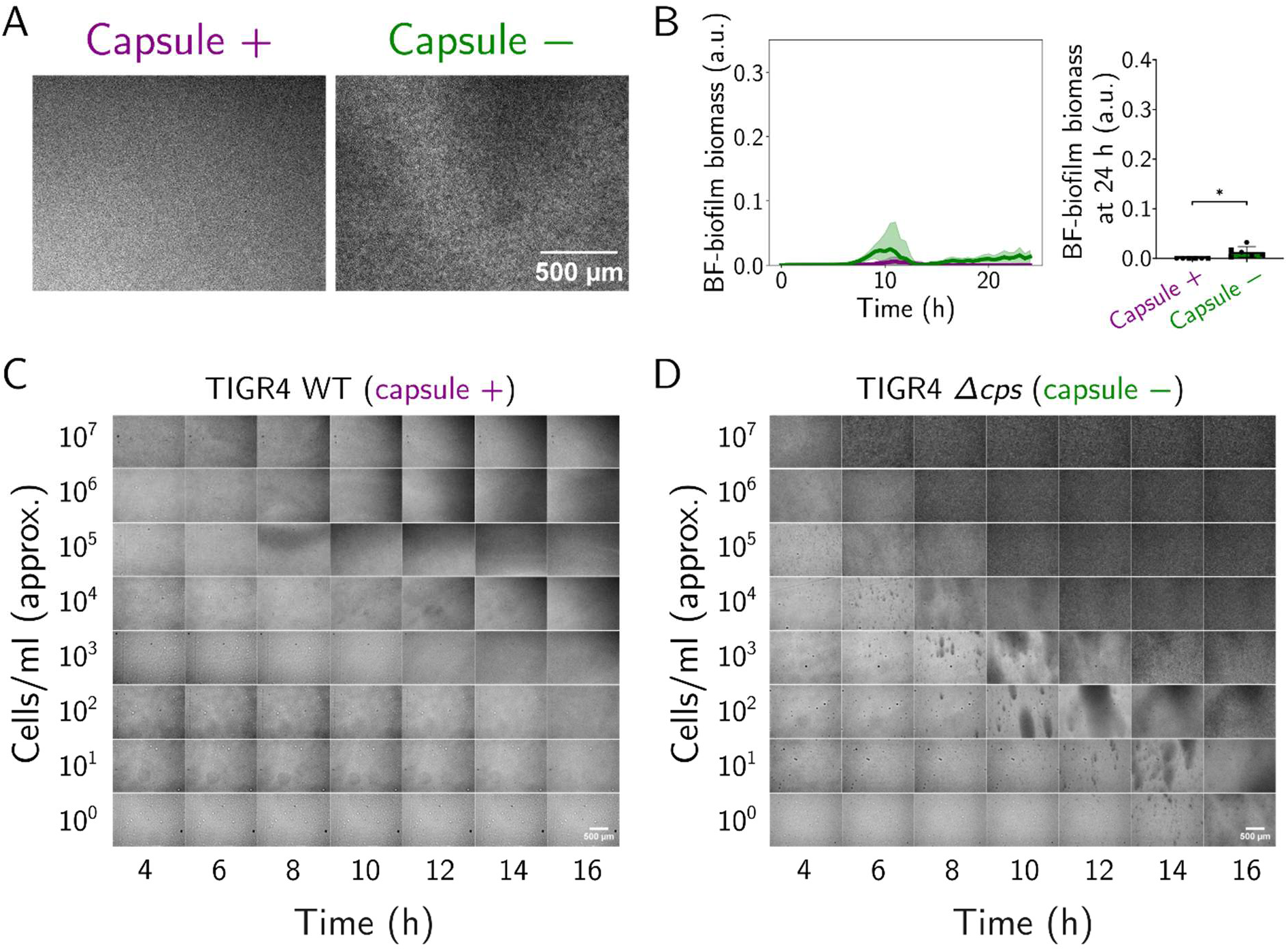
Model strain TIGR4 does not produce microcolonies, even when unencapsulated. The unencapsulated TIGR4 biofilms appeared thicker and more “grainy” in appearance at 24 hours post seeding compared to the encapsulated TIGR4, which is reflected as a small difference in microcolony biofilm biomass when quantified by LFAB. However, neither strain produces microcolony biofilms at any seeding density. **(A)** Brightfield microscopy images (4x magnification) of model strain TIGR4 WT (type 2 capsule; left) and isogenic unencapsulated (TIGR4 *Δcps*) mutant (right) at 24 hours post seeding, grown from ∼10^3^ cells/mL. **(B)** Time series plot (left) shows microcolony biofilm biomass for TIGR4 WT (purple) and *Δcps* (green) strains, quantified by LFAB from 0-24 hours post seeding at 30 min intervals. Line shows mean and shaded region shows standard deviations. Bar plot (right) shows the 24 hour timepoint. **(C)** Time series brightfield images of TIGR4 WT (encapsulated) at various initial cell seeding densities, showing highest seeding density at the top. **(D)** As in C, but for TIGR4 *Δcps* (unencapsulated). *N = 3* biological replicates with *n = 3* technical replicates each. Student’s two-tailed t-test; * *p < 0.05*. BF: brightfield. a.u.: arbitrary units. Scale bars are as indicated in each panel on the bottom right.

**Figure S6.**
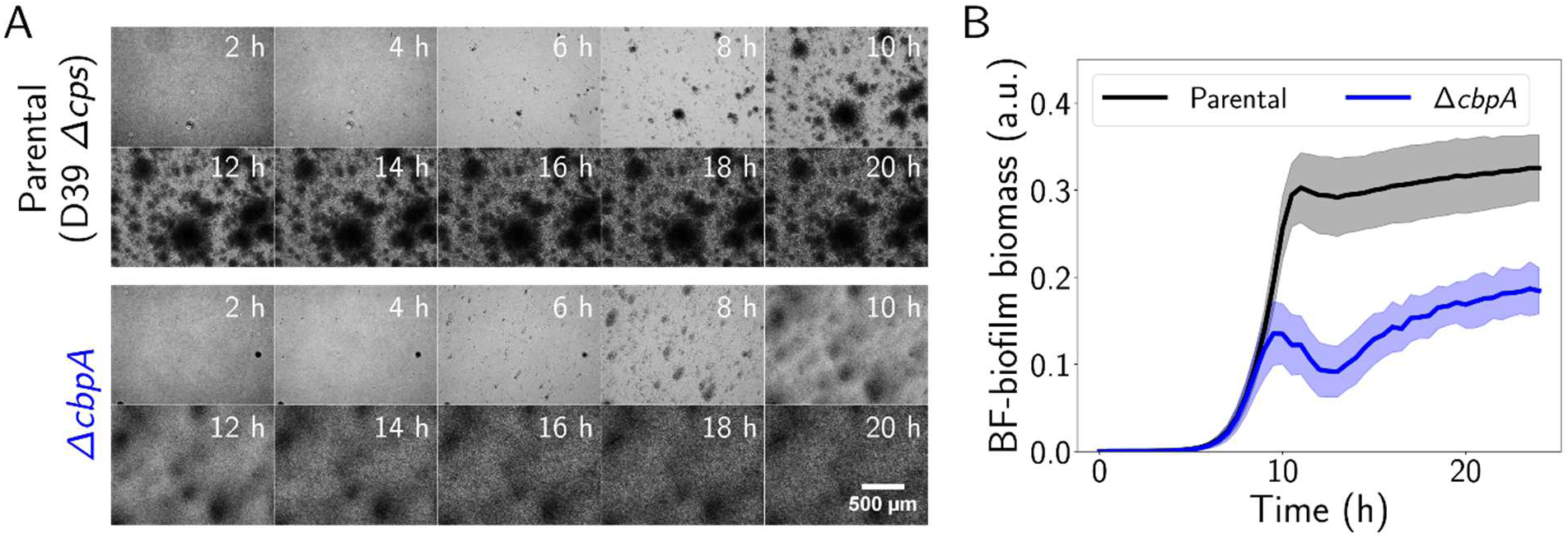
Longitudinal analysis of parental (D39*Δcps*) and *ΔcbpA* biofilm development. **(A)** Time series of biofilm development in the parental strain (D39*Δcps*; top) and *ΔcbpA* (bottom), showing images at indicated time points. Scale bar is the same for all images and is indicated on the bottom right. **(B)** LFAB quantification of the same strains, from 0-24 hours post seeding at 30 min intervals. Line shows mean and shaded region shows standard deviations. *N = 3* biological replicates with *n = 3* technical replicates each. BF: brightfield.

**Figure S7.**
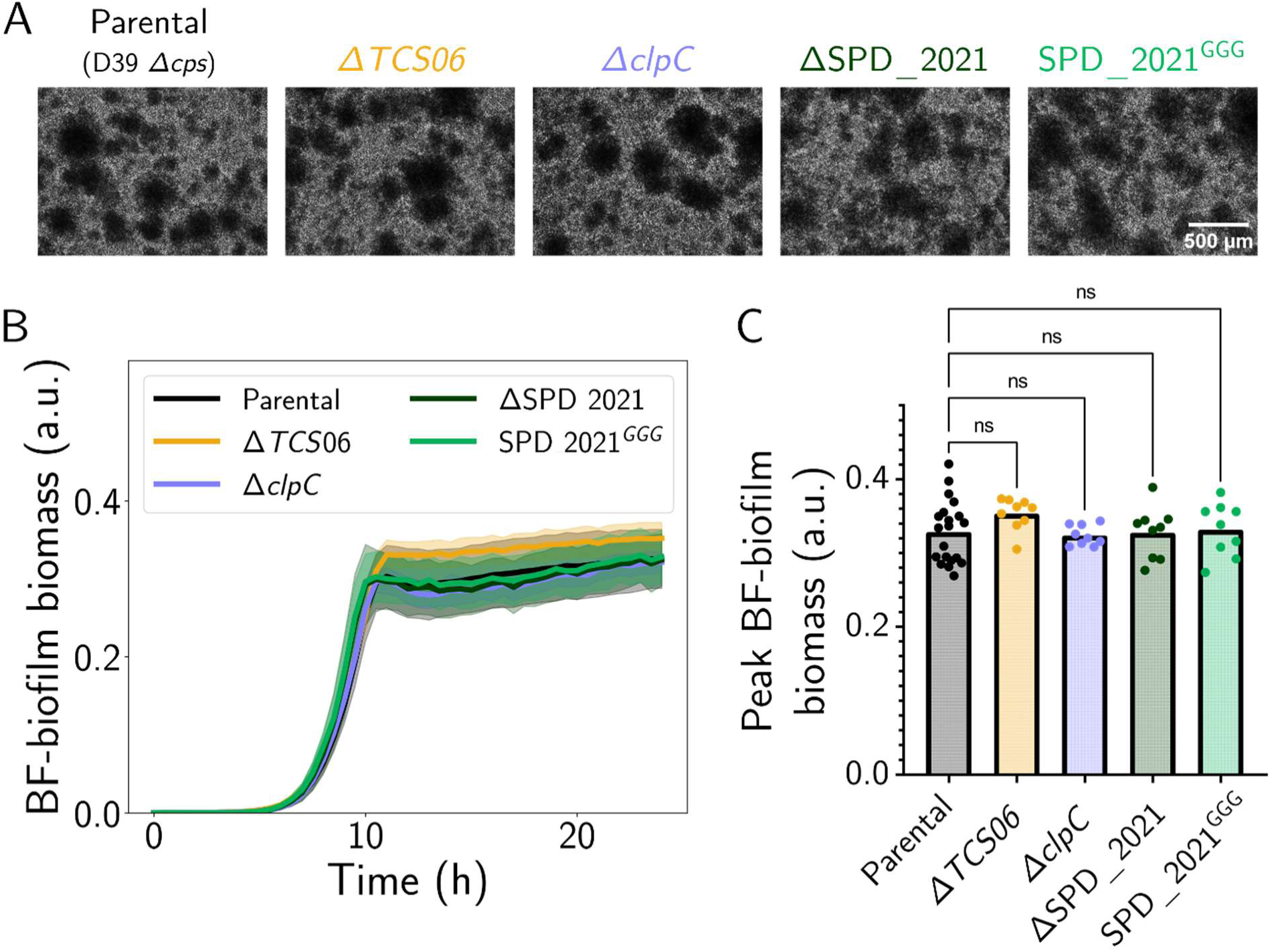
Deletion of *TCS06* (both *hk06* and *rr06*), *clpC*, or SPD_2021, as well as mutating the start codon of SPD_2021 has no effect on microcolony biofilms. *Our Tn-screen revealed some additional mutations in the* cbpA *genomic locus that phenocopied* ΔcbpA*. Specifically, insertion mutations in (i) the region upstream of* rr06 *in the hypothetical peptide SPD_2021, and (ii) in the* clp *protease. Here we show that deletion of these genes, as well as blocking translation of the predicted peptide, resembled the wild-type phenotype suggesting the biofilm defect may have been caused by polar effects of the transposon insertions.* (A) Brightfield biofilm images (4x magnification) of the parental strain (D39*Δcps*), *ΔTCS06*, *ΔclpC*, ΔSPD_2021 and SPD_2021^GGG^ (start codon mutated to ‘GGG’) at 24 hours post-seeding. **(B)** Time series of microcolony biofilm biomass for the above strains, as quantified by LFAB. **(C)** Peak microcolony biofilm biomass for each strain. For all strains, *N = 3-10* biological replicates, with *n = 3* technical replicates each. (A) Scale bars as shown. (B) Lines show the mean of all biological and technical replicates, and shaded areas show standard deviations. (C) Each data point represents the peak biomass of an individual time series. One way ANOVA, *p = 0.34* (not significant). Dunnett’s multiple comparisons test (comparing each strain to the parental); ns: not significant. BF: brightfield. a.u.: arbitrary units.

## Supplementary Tables

Supplementary Table S1: Descriptions of hits from transposon mutant screen, showing all mutants having microcolony biofilm biomass higher or lower than the parental (**Fig. 4C** shows the corresponding images)

Table provided as a separate Excel spreadsheet in Supplementary material: Supplementary_table_S1_Tn-screen_hits_details.xlsx

**Supplementary Table 2:**
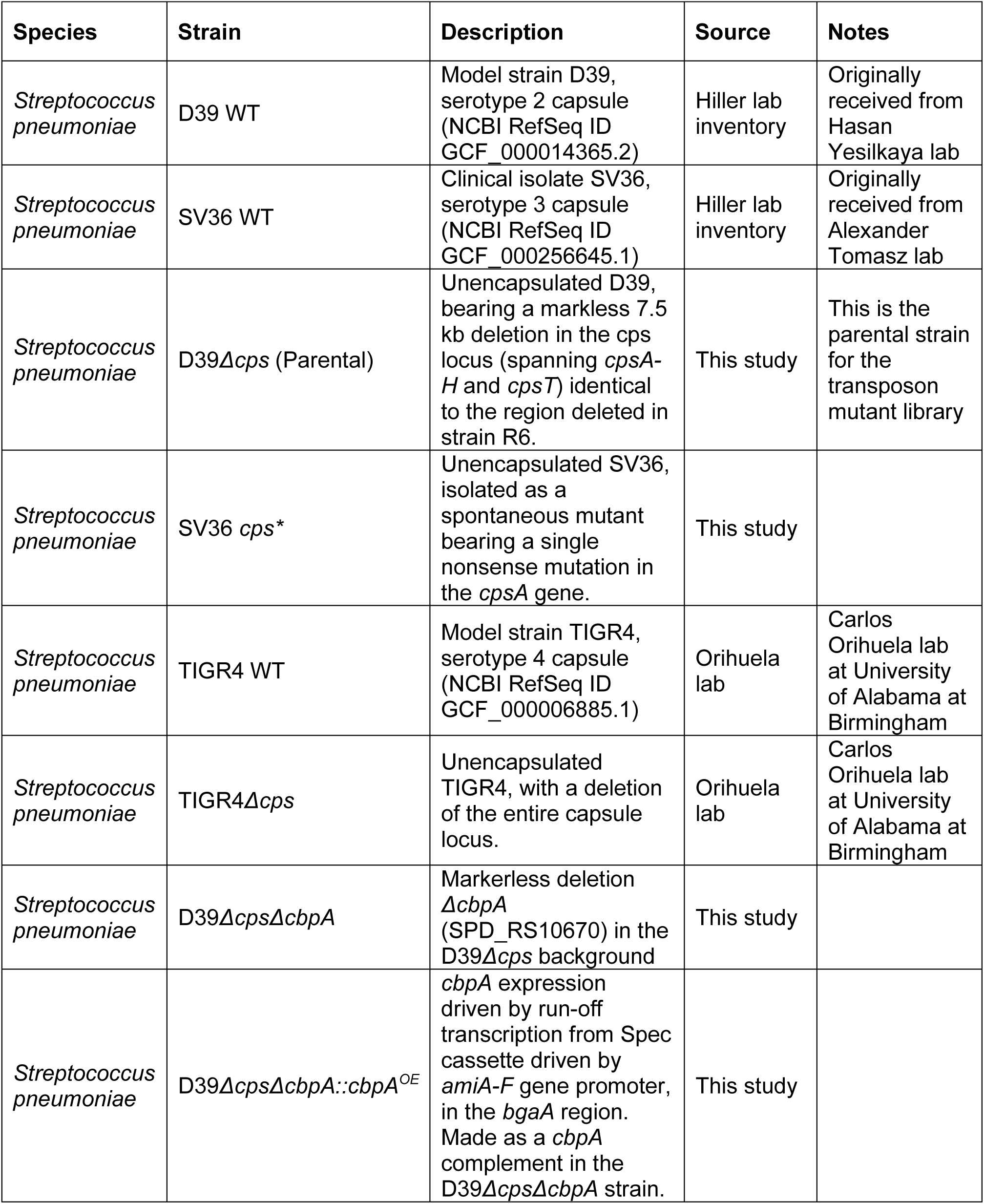
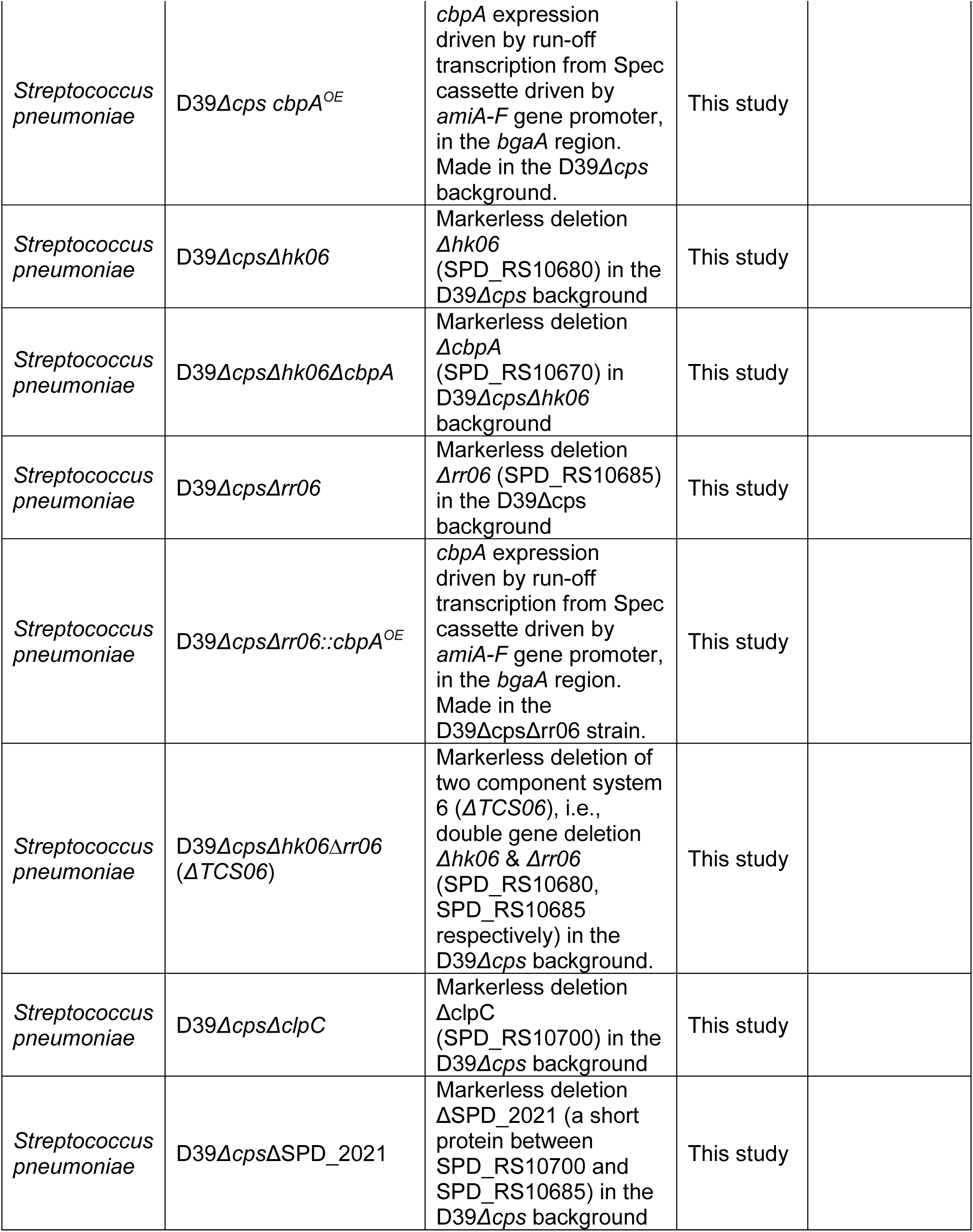
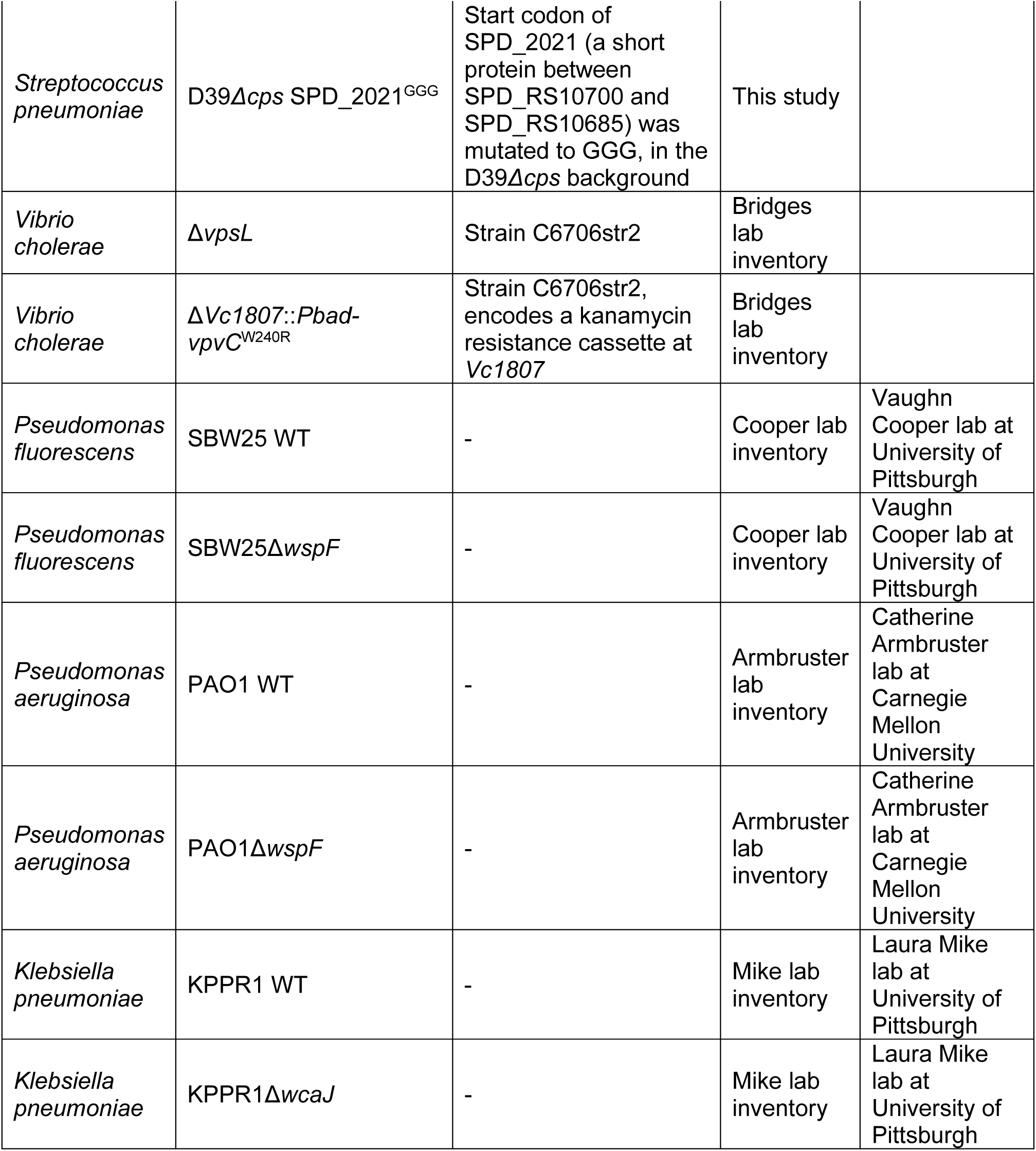
List of strains used in this study.

**Supplementary Table 3:**
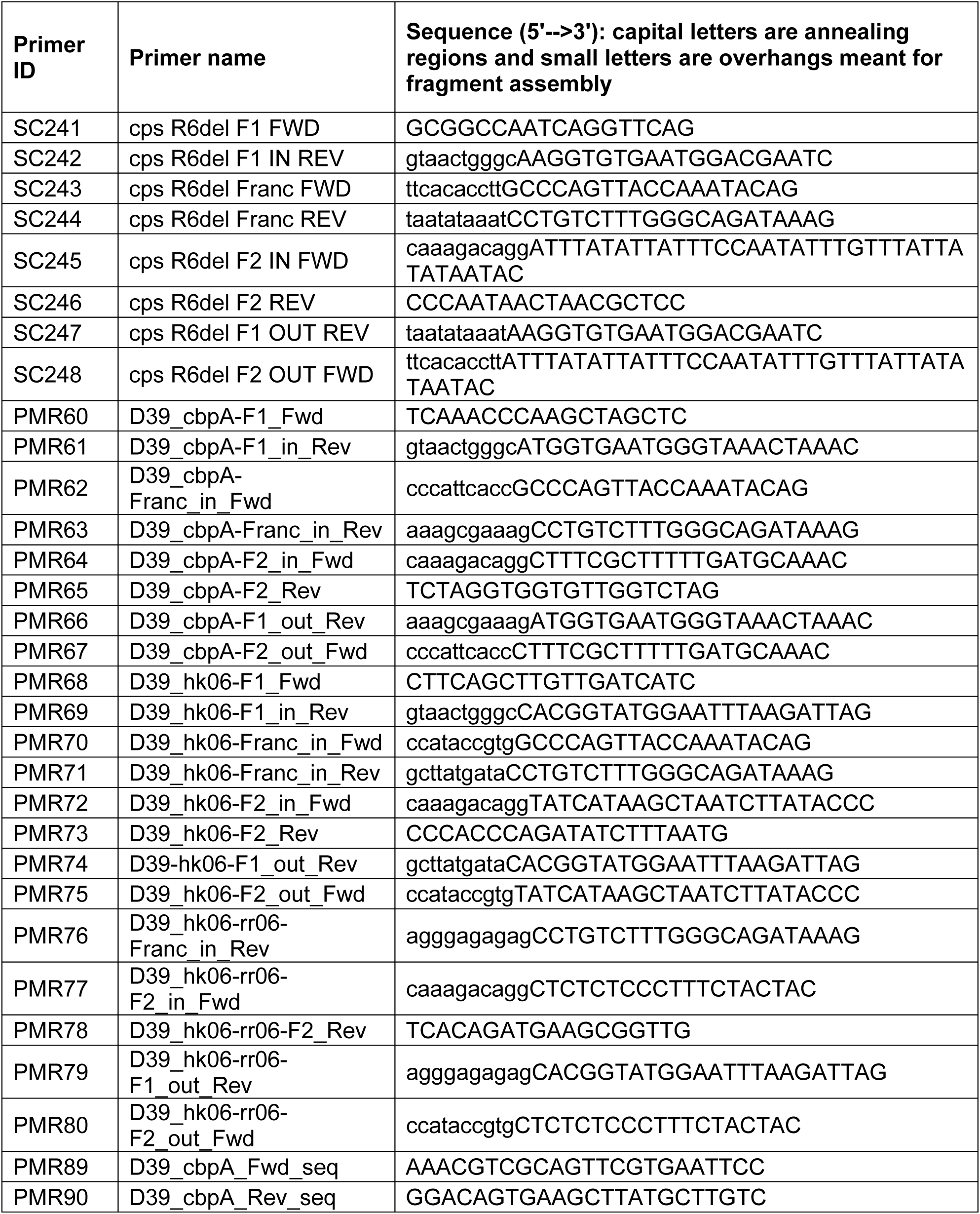
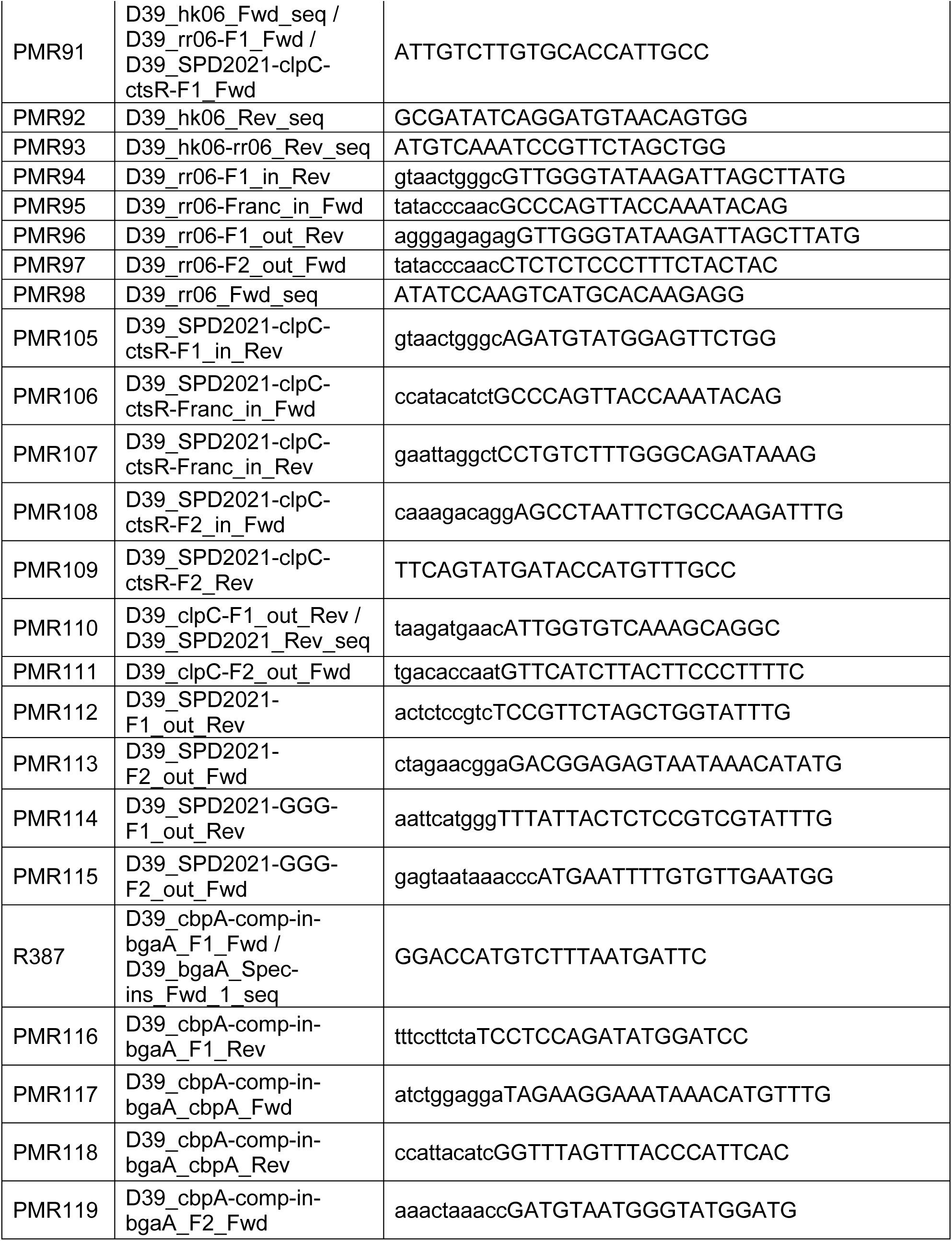
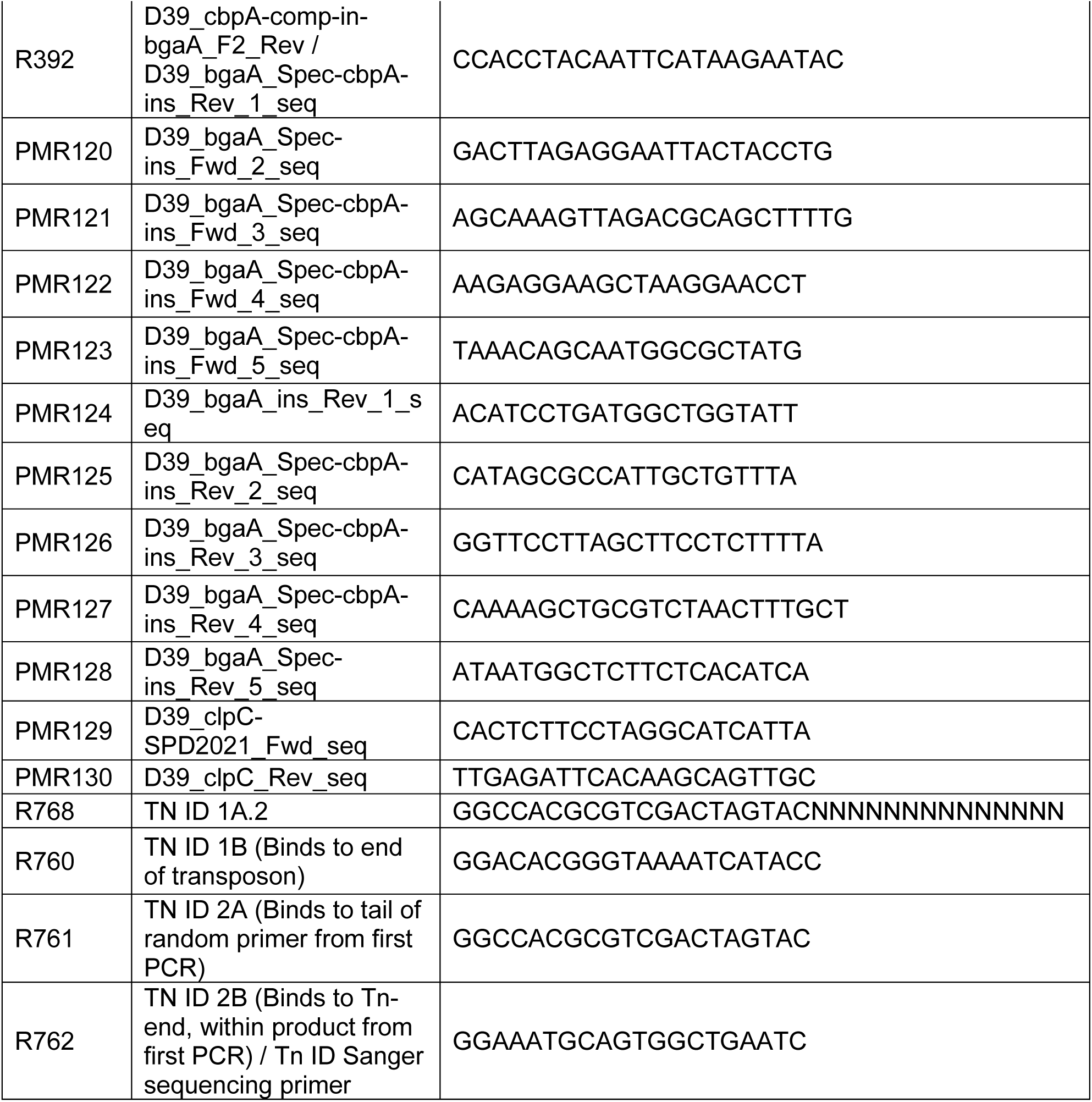
List of primers used in this study.

## References

1. Flemming, H.-C., et al. Biofilms: an emergent form of bacterial life. Nature Reviews Microbiology 14, 563–575 (2016).

2. Sauer, K., et al. The biofilm life cycle: expanding the conceptual model of biofilm formation. Nat Rev Microbiol 20, 608–620 (2022).

3. Yan, J. & Bassler, B. L. Surviving as a Community: Antibiotic Tolerance and Persistence in Bacterial Biofilms. Cell Host Microbe 26, 15–21 (2019).

4. Grari, O., et al. A comprehensive review on biofilm-associated infections: Mechanisms, diagnostic challenges, and innovative therapeutic strategies. The Microbe 8, 100436 (2025).

5. Iaconis, A., De Plano, L. M., Caccamo, A., Franco, D. & Conoci, S. Anti-Biofilm Strategies: A Focused Review on Innovative Approaches. Microorganisms 12, 639 (2024).

6. Roy, R., Tiwari, M., Donelli, G. & Tiwari, V. Strategies for combating bacterial biofilms: A focus on anti-biofilm agents and their mechanisms of action. Virulence 9, 522–554 (2017).

7. O’Toole, G. A. Microtiter Dish Biofilm Formation Assay. Journal of Visualized Experiments: JoVE (2011) doi:10.3791/2437.

8. Merritt, J. H., Kadouri, D. E. & O’Toole, G. A. Growing and Analyzing Static Biofilms. Current Protocols in Microbiology 22, 1B.1.1-1B.1.18 (2011).

9. Reichhardt, C. & Parsek, M. R. Confocal Laser Scanning Microscopy for Analysis of Pseudomonas aeruginosa Biofilm Architecture and Matrix Localization. Front Microbiol 10, 677 (2019).

10. Drescher, K., et al. Architectural transitions in Vibrio cholerae biofilms at single-cell resolution. Proc. Natl. Acad. Sci. U.S.A. 113, E2066–2072 (2016).

11. Qin, B., et al. Cell position fates and collective fountain flow in bacterial biofilms revealed by light-sheet microscopy. Science (2020) doi:10.1126/science.abb8501.

12. Yan, J., Sharo, A. G., Stone, H. A., Wingreen, N. S. & Bassler, B. L. Vibrio cholerae biofilm growth program and architecture revealed by single-cell live imaging. PNAS 113, E5337–E5343 (2016).

13. Prentice, J. A., Kasivisweswaran, S., Weerd, R. van de & Bridges, A. A. Biofilm dispersal patterns revealed using far-red fluorogenic probes. PLOS Biology 22, e3002928 (2024).

14. Berk, V., et al. Molecular Architecture and Assembly Principles of Vibrio cholerae Biofilms. Science 337, 236–239 (2012).

15. Bridges, A. A. & Bassler, B. L. The intragenus and interspecies quorum-sensing autoinducers exert distinct control over Vibrio cholerae biofilm formation and dispersal. PLoS Biol. 17, e3000429 (2019).

16. Bridges, A. A., Fei, C. & Bassler, B. L. Identification of signaling pathways, matrix-digestion enzymes, and motility components controlling Vibrio cholerae biofilm dispersal. Proc Natl Acad Sci U S A 117, 32639–32647 (2020).

17. Ikuta, K. S., et al. Global mortality associated with 33 bacterial pathogens in 2019: a systematic analysis for the Global Burden of Disease Study 2019. The Lancet 400, 2221–2248 (2022).

18. Chao, Y., Marks, L. R., Pettigrew, M. M. & Hakansson, A. P. Streptococcus pneumoniae biofilm formation and dispersion during colonization and disease. Front. Cell. Infect. Microbiol. 4, (2015).

19. Marks, L. R., Parameswaran, G. I. & Hakansson, A. P. Pneumococcal Interactions with Epithelial Cells Are Crucial for Optimal Biofilm Formation and Colonization In Vitro and In Vivo. Infection and Immunity (2012) doi:10.1128/iai.00488-12.

20. Gilley, R. P. & Orihuela, C. J. Pneumococci in biofilms are non-invasive: implications on nasopharyngeal colonization. Front. Cell. Infect. Microbiol. 4, (2014).

21. Gray, B. M., et al. Epidemiologic Studies of Streptococcus pneumoniae in Infants: Antibody Response to Nasopharyngeal Carriage of Types 3, 19, and 23.

22. de Lencastre, H. & Tomasz, A. From ecological reservoir to disease: the nasopharynx, day-care centres and drug-resistant clones of Streptococcus pneumoniae. J Antimicrob Chemother 50, 75–82 (2002).

23. Hall-Stoodley, L. et al. Direct Detection of Bacterial Biofilms on the Middle-Ear Mucosa of Children With Chronic Otitis Media.

24. Shenoy, A. T., et al. Streptococcus pneumoniae in the heart subvert the host response through biofilm-mediated resident macrophage killing. PLOS Pathogens 13, e1006582 (2017).

25. Brown, A. O., Millett, E. R. C., Quint, J. K. & Orihuela, C. J. Cardiotoxicity during Invasive Pneumococcal Disease. American Journal of Respiratory and Critical Care Medicine (2015) doi:10.1164/rccm.201411-1951PP.

26. Marks, L. R., Reddinger, R. M. & Hakansson, A. P. High Levels of Genetic Recombination during Nasopharyngeal Carriage and Biofilm Formation in Streptococcus pneumoniae. mBio (2012) doi:10.1128/mbio.00200-12.

27. Sanchez, C. J., et al. Streptococcus pneumoniae in Biofilms Are Unable to Cause Invasive Disease Due to Altered Virulence Determinant Production. PLOS ONE 6, e28738 (2011).

28. Hall-Stoodley, L., et al. Characterization of biofilm matrix, degradation by DNase treatment and evidence of capsule downregulation in Streptococcus pneumoniae clinical isolates. BMC Microbiology 8, 173 (2008).

29. Sanchez, C. J., et al. Biofilm and planktonic pneumococci demonstrate disparate immunoreactivity to human convalescent sera. BMC Microbiology 11, 245 (2011).

30. Kostyukova, N. N., Н, К. Н., Bekhalo, V. A. & А, Б. В. PNEUMOCOCCAL BIOFILMS AS A FORM OF PERSISTENCE: FORMATION, STRUCTURE, ROLE IN PATHOGENESIS, IMMUNE RESPONSE. Journal of microbiology, epidemiology and immunobiology 92, 55–62 (2015).

31. Muñoz-Elías, E. J., Marcano, J. & Camilli, A. Isolation of Streptococcus pneumoniae Biofilm Mutants and Their Characterization during Nasopharyngeal Colonization. Infection and Immunity 76, 5049–5061 (2008).

32. Rosenow, C., et al. Contribution of novel choline-binding proteins to adherence, colonization and immunogenicity of Streptococcus pneumoniae. Molecular Microbiology 25, 819–829 (1997).

33. Brooks-Walter, A., Briles, D. E. & Hollingshead, S. K. The pspC gene of Streptococcus pneumoniae encodes a polymorphic protein, PspC, which elicits cross-reactive antibodies to PspA and provides immunity to pneumococcal bacteremia. Infect Immun 67, 6533–6542 (1999).

34. Hammerschmidt, S., Talay, S. R., Brandtzaeg, P. & Chhatwal, G. S. SpsA, a novel pneumococcal surface protein with specific binding to secretory Immunoglobulin A and secretory component.

35. Janulczyk, R., Iannelli, F., Sjöholm, A. G., Pozzi, G. & Björck, L. Hic, a Novel Surface Protein of Streptococcus pneumoniae That Interferes with Complement Function*. Journal of Biological Chemistry 275, 37257–37263 (2000).

36. Standish, A. J., Stroeher, U. H. & Paton, J. C. The two-component signal transduction system RR06/HK06 regulates expression of cbpA in Streptococcus pneumoniae. Proceedings of the National Academy of Sciences 102, 7701–7706 (2005).

37. Standish, A. J., Stroeher, U. H. & Paton, J. C. The Pneumococcal Two-Component Signal Transduction System RR/HK06 Regulates CbpA and PspA by Two Distinct Mechanisms. Journal of Bacteriology (2007) doi:10.1128/jb.00335-07.

38. Ma, Z. & Zhang, J.-R. RR06 Activates Transcription of spr1996 and cbpA in Streptococcus pneumoniae. Journal of Bacteriology (2007) doi:10.1128/jb.01429-06.

39. Moscoso, M., García, E. & López, R. Biofilm Formation by Streptococcus pneumoniae: Role of Choline, Extracellular DNA, and Capsular Polysaccharide in Microbial Accretion. Journal of Bacteriology (2006) doi:10.1128/jb.00673-06.

40. Fong, J. C. N., Syed, K. A., Klose, K. E. & Yildiz, F. H. Role of Vibrio polysaccharide (vps) genes in VPS production, biofilm formation and Vibrio cholerae pathogenesis. Microbiology (Reading, Engl.) 156, 2757–2769 (2010).

41. Bridges, A. A., Prentice, J. A., Fei, C., Wingreen, N. S. & Bassler, B. L. Quantitative input– output dynamics of a c-di-GMP signal transduction cascade in Vibrio cholerae. PLOS Biology 20, e3001585 (2022).

42. Domenech, M., García, E. & Moscoso, M. Versatility of the capsular genes during biofilm formation by Streptococcus pneumoniae. Environmental Microbiology 11, 2542–2555 (2009).

43. Qin, L., Kida, Y., Imamura, Y., Kuwano, K. & Watanabe, H. Impaired capsular polysaccharide is relevant to enhanced biofilm formation and lower virulence in Streptococcus pneumoniae. J Infect Chemother 19, 261–271 (2013).

44. Hammerschmidt, S., et al. Illustration of Pneumococcal Polysaccharide Capsule during Adherence and Invasion of Epithelial Cells. Infection and Immunity 73, 4653–4667 (2005).

45. Parker, D., et al. The NanA Neuraminidase of Streptococcus pneumoniae Is Involved in Biofilm Formation. Infect Immun 77, 3722–3730 (2009).

46. Blanchette-Cain, K., et al. Streptococcus pneumoniae Biofilm Formation Is Strain Dependent, Multifactorial, and Associated with Reduced Invasiveness and Immunoreactivity during Colonization. mBio 4, 10.1128/mbio.00745-13 (2013).

47. Wei, H. & Håvarstein, L. S. Fratricide is essential for efficient gene transfer between pneumococci in biofilms. Appl Environ Microbiol 78, 5897–5905 (2012).

48. Vidal, J. E., Ludewick, H. P., Kunkel, R. M., Zähner, D. & Klugman, K. P. The LuxS-dependent quorum-sensing system regulates early biofilm formation by Streptococcus pneumoniae strain D39. Infect Immun 79, 4050–4060 (2011).

49. Janssen, A. B., et al. PneumoBrowse 2: an integrated visual platform for curated genome annotation and multiomics data analysis of Streptococcus pneumoniae. Nucleic Acids Res 53, D839–D851 (2025).

50. Aprianto, R., Slager, J., Holsappel, S. & Veening, J.-W. High-resolution analysis of the pneumococcal transcriptome under a wide range of infection-relevant conditions. Nucleic Acids Res 46, 9990–10006 (2018).

51. Slager, J., Aprianto, R. & Veening, J.-W. Deep genome annotation of the opportunistic human pathogen Streptococcus pneumoniae D39. Nucleic Acids Res 46, 9971–9989 (2018).

52. Buckwalter, C. M. & King, S. J. Pneumococcal carbohydrate transport: food for thought. Trends Microbiol 20, 517–522 (2012).

53. Complete Genome Sequence of a Virulent Isolate of Streptococcus pneumoniae | Science. https://www.science.org/doi/10.1126/science.1061217.

54. Streptococcus pneumoniae Capsular Polysaccharide | Microbiology Spectrum. https://journals.asm.org/doi/10.1128/microbiolspec.gpp3-0019-2018?url_ver=Z39.88-2003&rfr_id=ori%3Arid%3Acrossref.org&rfr_dat=cr_pub++0pubmed.

55. Vollmer, W., Massidda, O. & Tomasz, A. The Cell Wall of Streptococcus pneumoniae. Microbiology Spectrum 7, 10.1128/microbiolspec.gpp3-0018–2018 (2019).

56. Filipe, S. R., Pinho, M. G. & Tomasz, A. Characterization of the murMN Operon Involved in the Synthesis of Branched Peptidoglycan Peptides in Streptococcus pneumoniae *. Journal of Biological Chemistry 275, 27768–27774 (2000).

57. Filipe, S. R. & Tomasz, A. Inhibition of the expression of penicillin resistance in Streptococcus pneumoniae by inactivation of cell wall muropeptide branching genes. Proceedings of the National Academy of Sciences 97, 4891–4896 (2000).

58. Filipe, S. R., Severina, E. & Tomasz, A. The Role of murMN Operon in Penicillin Resistance and Antibiotic Tolerance of Streptococcus pneumoniae. Microbial Drug Resistance 7, 303– 316 (2001).

59. Filipe, S. R., Severina, E. & Tomasz, A. The murMN operon: A functional link between antibiotic resistance and antibiotic tolerance in Streptococcus pneumoniae. Proceedings of the National Academy of Sciences 99, 1550–1555 (2002).

60. A molecular link between cell wall biosynthesis, translation fidelity, and stringent response in Streptococcus pneumoniae | PNAS. https://www.pnas.org/doi/10.1073/pnas.2018089118.

61. Charpentier, E., Novak, R. & Tuomanen, E. Regulation of growth inhibition at high temperature, autolysis, transformation and adherence in Streptococcus pneumoniae by ClpC. Molecular Microbiology 37, 717–726 (2000).

62. Iannelli, F., Oggioni, M. R. & Pozzi, G. Allelic variation in the highly polymorphic locus pspC of Streptococcus pneumoniae. Gene 284, 63–71 (2002).

63. van der Maten, E., et al. Streptococcus pneumoniae PspC Subgroup Prevalence in Invasive Disease and Differences in Contribution to Complement Evasion. Infection and Immunity 86, 10.1128/iai.00010-18 (2018).

64. Maestro, B. & Sanz, J. M. Choline Binding Proteins from Streptococcus pneumoniae: A Dual Role as Enzybiotics and Targets for the Design of New Antimicrobials. Antibiotics (Basel) 5, 21 (2016).

65. Sinha, D., Frick, J. P., Clemons, K., Winkler, M. E. & De Lay, N. R. Pivotal Roles for Ribonucleases in Streptococcus pneumoniae Pathogenesis. mBio 12, e02385–21.

66. Berry, A. M., Lock, R. A. & Paton, J. C. Cloning and characterization of nanB, a second Streptococcus pneumoniae neuraminidase gene, and purification of the NanB enzyme from recombinant Escherichia coli. Journal of Bacteriology 178, 4854–4860 (1996).

67. Jensch, I., et al. PavB is a surface-exposed adhesin of Streptococcus pneumoniae contributing to nasopharyngeal colonization and airways infections. Molecular Microbiology 77, 22–43 (2010).

68. Cámara, M., Boulnois, G. J., Andrew, P. W. & Mitchell, T. J. A neuraminidase from Streptococcus pneumoniae has the features of a surface protein. Infection and Immunity 62, 3688–3695 (1994).

69. Zhao, G., Meier, T. I., Hoskins, J. & McAllister, K. A. Identification and Characterization of the Penicillin-Binding Protein 2a of Streptococcus pneumoniae and Its Possible Role in Resistance to β-Lactam Antibiotics. Antimicrob Agents Chemother 44, 1745–1748 (2000).

70. A Functional dlt Operon, Encoding Proteins Required for Incorporation of d-Alanine in Teichoic Acids in Gram-Positive Bacteria, Confers Resistance to Cationic Antimicrobial Peptides in Streptococcus pneumoniae | Journal of Bacteriology. https://journals.asm.org/doi/10.1128/jb.00336-06?url_ver=Z39.88-2003&rfr_id=ori%3Arid%3Acrossref.org&rfr_dat=cr_pub++0pubmed.

71. De Saizieu, A., et al. Microarray-based identification of a novel Streptococcus pneumoniae regulon controlled by an autoinduced peptide. J. Bacteriol. 182, 4696–4703 (2000).

72. Dawid, S., Roche, A. M. & Weiser, J. N. The blp bacteriocins of Streptococcus pneumoniae mediate intraspecies competition both in vitro and in vivo. Infect. Immun. 75, 443–451 (2007).

73. Shlla, B., et al. The Rgg1518 transcriptional regulator is a necessary facet of sugar metabolism and virulence in Streptococcus pneumoniae. Molecular Microbiology 116, 996–1008 (2021).

74. Bergé, M. J., et al. Midcell Recruitment of the DNA Uptake and Virulence Nuclease, EndA, for Pneumococcal Transformation. PLoS Pathog 9, e1003596 (2013).

75. Search for Genes Essential for Pneumococcal Transformation: the RadA DNA Repair Protein Plays a Role in Genomic Recombination of Donor DNA | Journal of Bacteriology. https://journals.asm.org/doi/10.1128/jb.00573-07?url_ver=Z39.88-2003&rfr_id=ori%3Arid%3Acrossref.org&rfr_dat=cr_pub++0pubmed.

76. Afzal, M., Manzoor, I., Kuipers, O. P. & Shafeeq, S. Cysteine-Mediated Gene Expression and Characterization of the CmbR Regulon in Streptococcus pneumoniae. Front Microbiol 7, 1929 (2016).

77. Guizar-Sicairos, M., Thurman, S. T. & Fienup, J. R. Efficient subpixel image registration algorithms. Opt. Lett, OL 33, 156–158 (2008).

78. Mueller Brown, K., et al. Peptide maturation molecules act as molecular gatekeepers to coordinate cell-cell communication in Streptococcus pneumoniae. Cell Rep 43, 114432 (2024).

79. Smith, A. P., et al. Dynamic Pneumococcal Genetic Adaptations Support Bacterial Growth and Inflammation during Coinfection with Influenza. Infection and Immunity 89, (2021).

80. Domenech, M. & García, E. The N-Acetylglucosaminidase LytB of Streptococcus pneumoniae Is Involved in the Structure and Formation of Biofilms. Applied and Environmental Microbiology 86, e00280–20 (2020).

81. Rico-Lastres, P., et al. Substrate recognition and catalysis by LytB, a pneumococcal peptidoglycan hydrolase involved in virulence. Sci Rep 5, 16198 (2015).

82. De Las Rivas, B., García, J. L., López, R. & García, P. Purification and polar localization of pneumococcal LytB, a putative endo-beta-N-acetylglucosaminidase: the chain-dispersing murein hydrolase. J Bacteriol 184, 4988–5000 (2002).

83. García, P., González, M. P., García, E., López, R. & García, J. L. LytB, a novel pneumococcal murein hydrolase essential for cell separation. Molecular Microbiology 31, 1275–1277 (1999).

84. Domenech, A., Slager, J. & Veening, J.-W. Antibiotic-Induced Cell Chaining Triggers Pneumococcal Competence by Reshaping Quorum Sensing to Autocrine-Like Signaling. Cell Reports 25, 2390–2400.e3 (2018).

85. Aggarwal, S. D., et al. Function of BriC peptide in the pneumococcal competence and virulence portfolio. PLOS Pathogens 14, e1007328 (2018).

86. Aggarwal, S. D., et al. BlpC-mediated selfish program leads to rapid loss of Streptococcus pneumoniae clonal diversity during infection. Cell Host Microbe 31, 124–134.e5 (2023).

87. Hobbs, J. K., Pluvinage, B. & Boraston, A. B. Glycan-metabolizing enzymes in microbe–host interactions: the Streptococcus pneumoniae paradigm. FEBS Letters 592, 3865–3897 (2018).

88. Uchiyama, S., et al. The surface-anchored NanA protein promotes pneumococcal brain endothelial cell invasion. J Exp Med 206, 1845–1852 (2009).

